# Moving object generates topographic responses in the majority of entorhinal neurons

**DOI:** 10.64898/2026.10.05.756590

**Authors:** Chinmay S. Purandare, Severine Durand, Ben Hardcastle, Ryan Gillis, Mikayla Carlson, Hannah Belski, Carter Peene, Ahad Bawany, Siddharth Jha, Avalon Amaya, Vivian Ha, Warren Han, Ben Ouellette, Tye Johnson, Robyn Naidoo, Katrina Nguyen, Lucas Suarez, Josh Wilkes, Ahrial Young, Shiella Caldejon, Jackie Swapp, Ali Williford, Peter A. Groblewski, Jérôme A. Lecoq, Mayank R. Mehta

## Abstract

Vision is crucial in generating a complex array of responses in rodents during spatial exploration and in humans during memory tasks. These tasks involve other variables too, e.g. multisensory cues in rodents and complex memories in humans. Further, there are major differences between the medial and lateral entorhinal cortical (MEC and LEC) functional cell types. To understand the contribution of purely visual information to entorhinal responses, we presented a simple, moving vertical bar of light to head-fixed mice, without any task or reward. Surprisingly, 66% (2391/3601) of the MEC broad spiking neurons were significantly modulated by the bar position on the screen. Visual tuning in MEC had a pronounced dorsoventral topography –the tuning strength dropped from 79% in the dorsal most quartile to 40% in the ventral most quartile. Unlike major differences in spatial and other codes between MEC and LEC during free foraging experiments, LEC cells showed very similar visual tuning (392/606, 65%), including the dorso-ventral gradient in LEC5. Additionally, the majority of putative interneurons, from MEC as well as LEC, showed similar, widespread visual tuning, unlike spatial tasks. Neurons also encoded the object motion direction (naso-temporal vs. temporo-nasal visual motion) with changes in mean rate, tuning strength and/or tuning location. Such directional nature of visual tuning indicates significant history-dependence or sequence encoding. Consistently, replacing the continuous motion by randomly flashing bar of light abolished bar-tuning in two-thirds of continuous bar-tuned cells and substantially altered tuning in the rest. These results reveal the contribution of purely visual inputs in generating anatomically organized responses in the majority of entorhinal neurons. The prevalence and similarity of visual tuning across all layers of MEC as well as LEC exceeds that of other known functional cell types (like grid cells in MEC or egocentric object tuning/time-cells in LEC) during spatial navigation tasks and could be foundational in understanding entorhinal complexity.

## Introduction

The rodent entorhinal cortex has been studied extensively in the context of spatial navigation, leading to many seminal discoveries like the grid cells(1–5), border cells(6,7), speed cells(4,8), spatial cells(9,10), object vector(11,12) cells, cue cells(13–15), time(16) and head direction(4) cells. Most of these studies have focused on the medial subdivision of the entorhinal cortex (MEC), considered to be a part of the “where” pathway(17–19). The exact prevalence of these neural codes are debated(9,10).

In contrast to MEC, the lateral entorhinal cortex (LEC) is considered to be a part of the “what” pathway(17,18,20), and LEC neurons show much weaker spatial selectivity than MEC(20–22). Instead, they are thought to be governed strongly by multisensory cues(23), especially olfactory stimuli(24–28). Inclusion of multisensory cues, like objects in the maze, can improve LEC selectivity(29), but in the egocentric, not allocentric space(30). Further, some LEC cells encode the elapsed time(16). Most of these important functional cell types are represented by less than a third of active LEC cells, typically in some but not all sub-layers of LEC. Stimuli that generate selective responses in a majority of LEC neurons have remained elusive. Moreover, many of these studies involve specific, spatial or non-spatial tasks with distinct types of locomotion to obtain rewards, which also affects entorhinal encoding(31). Neurons upstream of the entorhinal cortex, e.g. the postrhinal cortex show significant tuning to visual stimuli during self-motion(32) or reward association tasks(33), and consistently receive visual information encoding passively viewed visual motion(34). Parietal cortex, another upstream input for the entorhinal cortex, encodes visual information(35,36). Further upstream, neurons from the primary sensory visual areas encode purely visual information, without any locomotion, rewards or tasks(37–39).

A subset of primate entorhinal neurons have grid-like firing patterns encoding visual space(40). Further, the majority of primate entorhinal cells encode gaze position on a screen (41–44), and are implicated in visual cue dependent memory tasks(45,46). In humans, damage to entorhinal cortex causes profound deficits in not only spatial(47,48) but non-spatial memories too, experienced by an immobile subject, e.g. object-identity or location on a screen(49). Consistently, human entorhinal neurons encode visually evoked responses to events, faces and objects(50–52).

However, purely visually evoked responses in the entorhinal cortex of immobile subjects without task or memory requirements are not reported in any species, even though these are commonly found in the upstream visual cortex in several species. Here we report such responses, which can form the basis of more complex representations to mediate a variety of tasks. Moreover, the visual responses we report here are strongly dependent on the sequential or continuous nature of the stimulus, which could be the basis for the role of entorhinal-hippocampal system in sequence encoding(53–56) during different tasks.

## Results

Nine mice (7 females) were head-fixed next to a large screen and placed on a running wheel (Fig. 1A). They were habituated to viewing simple stimuli, e.g. moving bars of lights (henceforth referred to as bars, for brevity) on the screen of an experimental rig. There were no task demands, self-motion requirements or rewards but the animals were free to run on a slanted running wheel. The bars were presented monocularly and their range of movements spanned most of the right hemifield (120°, Fig 1A). Neural signals were measured from the left hemisphere (Fig. 1B) using Neuropixel probes in MEC and/or LEC (Fig. 1C). The anatomical locations of the recorded neurons was determined post-hoc by registering the electrode locations with the Allen brain atlas(57) (see *Methods*).

**Figure 1.**
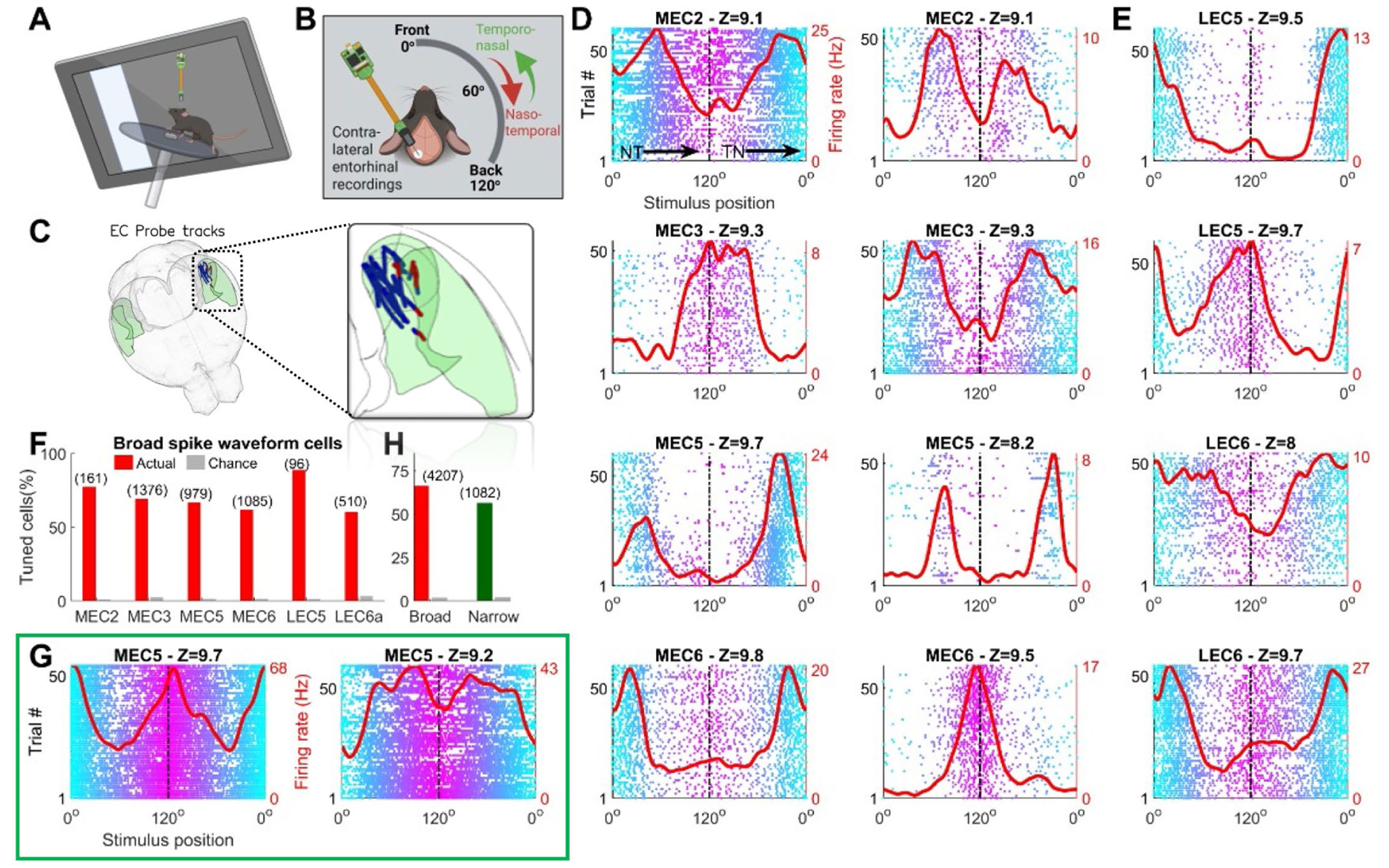
Medial and lateral entorhinal neurons encode the position of a moving bar of light. **(A)** Illustration of the setup showing a mouse (head-fixed) running on a wheel, while visual stimuli are shown on the right side. Neuropixels (version 1.0) probe(s) simultaneously recorded brain activity from the entorhinal cortex. **(B)** Illustration of the setup in a top view, depicting visual stimuli presentation on the right side, with neural recordings in the contralateral entorhinal cortex. Stimuli moved in naso-temporal (NT, green arrow) as well as temporo-nasal directions (red arrow). Although stimuli were presented on a flat screen, spherical warping was deployed to account for the close-viewing angle, allowing for presentation locations from the front (0°) to the back (120°) of the hemi-field as shown. The most nasal extent of the screen, set at 0° in convention, reached 7° on the left side. **(C)** Probe tracks through entorhinal cortex across all animals. Blue dots depict units in MEC, red in LEC. **(D)** 8 example cells from the medial entorhinal cortex (MEC) showing tuned responses for distinct locations of the bar of light. Red trace depicts mean response, cyan-magenta dots indicate spikes in individual trials, cyan corresponding to nasal locations, and magenta corresponding to temporal locations on the screen. Sublayer of recording is also indicated on the top. **(E)** Same as (D), for lateral entorhinal cortex (LEC). The first half of each response (0°-120°, front to back) corresponds with naso-temporal motion, 2^nd^ half (120°-0°, back-front) corresponds to temporo-nasal response. **(F)** Bar chart denoting fraction of cells across broad spiking units for 6 sublayers of EC (red-actual, gray-chance level). Cell count is indicated in parenthesis above. **(G)** Similar to (D), two cells (from MEC5) with high firing rate and narrow spike waveform (i.e., putative interneurons) with significant tuning for the bar position. **(H)** Bar chart denoting fraction of cells found to be visually tuned, separately for broad (n=4207), and narrow (1082) spike waveform cells. Visual tuning was significantly more prevalent in broad (66.2%), compared to narrow (55.1%) spike waveform units (KS-test, here and throughout, unless otherwise mentioned, *p*=1.3×10^−15^).

### Majority of entorhinal cells’ activity is modulated by a moving bar of light

Many putatively excitatory neurons (with broad spike waveforms) in MEC (Fig. 1D) and LEC (Fig. 1E) showed clear, succinct and reliable firing preference for a variety of bar locations. Simultaneously recorded EC cells were tuned for diverse positions of the bar, arguing against their non-specific noise related origins (fig. S1).

We next examined the prevalence of visual tuning in neurons across the 6 subregions of entorhinal cortex (MEC layers 2, 3, 5 and 6, and LEC layers 5 and 6). More than 50% of broad spiking cells across all parts of EC were significantly modulated (see *Methods* for quantification metrics) by the bar position, with the largest prevalence in LEC5 (88.5%), and the least in LEC6 (60.2%) (Fig. 1F). The mean firing rates across brain regions were qualitatively similar (3.9Hz-LEC6 to 5.3Hz-MEC3) but the depth of modulation was larger in LEC5 (2.3 vs ∼1.1-1.5 in other brain regions) (fig. S2). Thus, these differences in tuning prevalence across areas were not due to differences in mean firing rates, but due to stronger modulation.

Entorhinal spatial selectivity has been typically described in excitatory cells(1,2), which have broad spike waveforms and lower mean firing rates. On the other hand, putative inhibitory cells typically have narrow waveforms and higher mean firing rates (fig. S2). Given the widespread modulation in broad spiking cells, we wondered if the narrow spiking cells also had visual receptive fields. To our surprise, many narrow spiking cells were modulated by bar position (Fig. 1G). Across 1082 narrow spiking cells, 56.1% had significant tuning, which was comparable but slightly lower than that for broad spiking cells (Fig. 1H). Imposition of further restrictions based on complex spiking index and mean firing rates did not qualitatively change the prevalence of tuning (see *Methods*, and fig. S2). Thus, visual tuning in EC is robust and widespread, present in a majority of excitatory as well as inhibitory neurons.

### Dorsoventral gradient of entorhinal visual tuning

MEC grid cells show topographic organization, with increasing receptive field size as a function of the dorso-ventral depth(1,58). Several mechanisms have been proposed to explain this(59–63). Hence, we investigated the topographic organization of visual tuning. We first focused on MEC3, whose head-direction(59) as well as grid-cell tuning(4,58,61) in freely behaving rodents is dorso-ventrally graded. Visual tuning in MEC3 was far more prevalent in the dorsal than the ventral parts (Fig. 2A). The tuning prevalence showed a substantial reduction beyond the depth of about ∼3800um (Fig. 2B). This was not due to nonspecific factors such as changes in the mean firing rates (fig. S3). Similar topographic organization was also seen for MEC5 and MEC6 (Fig. 2B), contrary to previous reports claiming the lack of such gradient in these sub-layers for head direction selectivity during free-foraging(59). Dorso-ventral gradient was also seen in MEC2 (fig. S3). In fact, similar dorso-ventral gradient was found in all MEC areas, and as a result, also when combining all broad spiking MEC cells (Fig. 2C). Further, even the narrow spiking cells too showed a similar gradient (Fig. 2D). When combined across cell-classes and sub-layers, visual tuning in dorsal-most MEC was four times as prevalent as ventral-most EC (Fig. 2E). Taken together, MEC cells showed robust, pan-layer gradient in strength of tuning with stronger and more prevalent tuning in the dorsal, compared to the ventral parts. Interestingly, dorsal LEC5 too had significantly greater visual tuning than ventral (fig. S3). LEC6 also showed a similar trend (but not significant).

**Figure 2.**
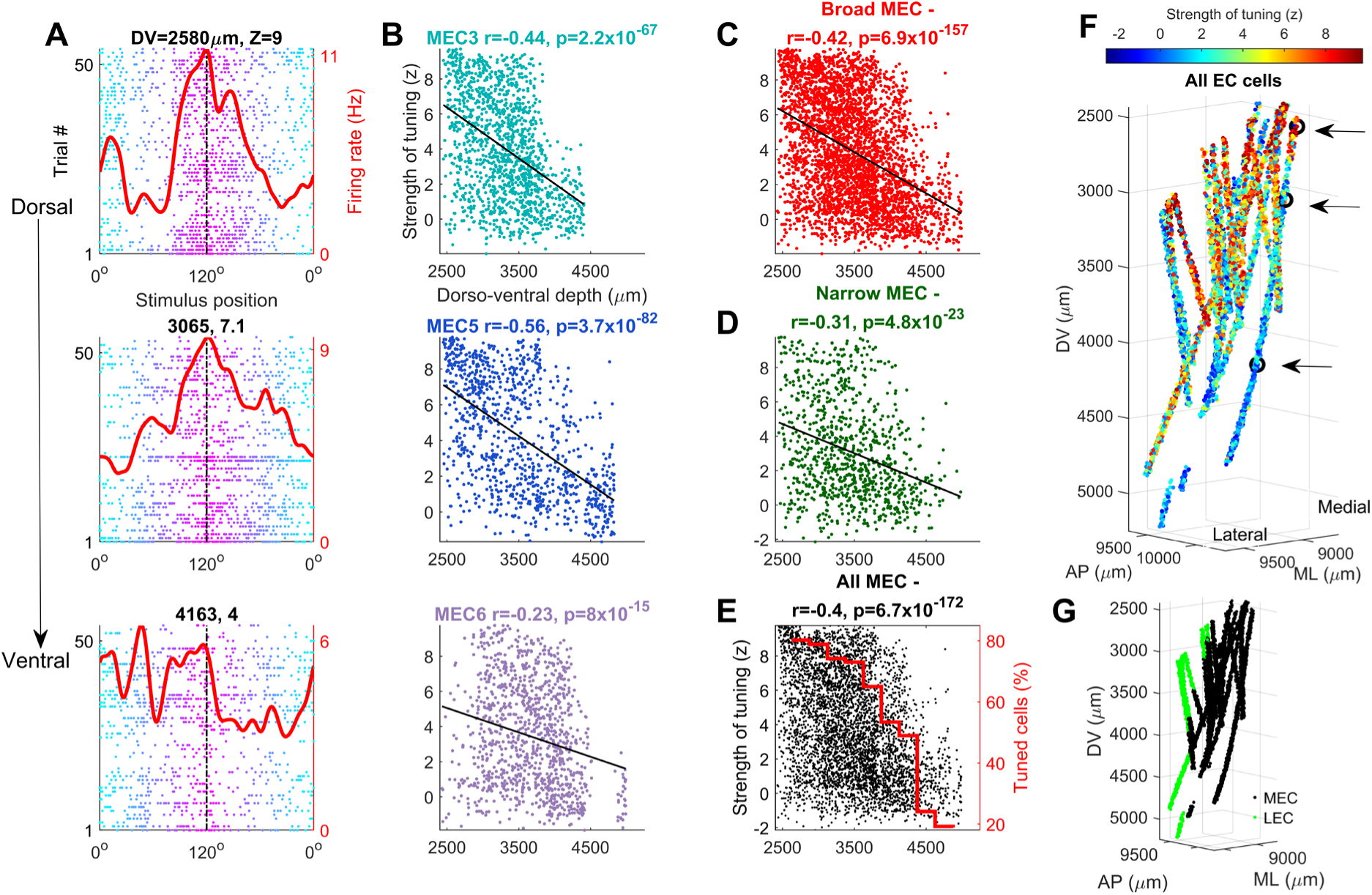
Dorso-ventral gradient of visual tuning in EC. **(A)** 3 simultaneously recorded MEC3 cells from the same *Neuropixels* shank arranged in the increasing order of their anatomical depth. Note-strength of tuning (indicated above) decreases for more ventral cells. **(B)** Scatter plot between the dorso-ventral location of a unit and its strength of tuning. MEC3 as well as 5 and 6 cells had significantly better visual tuning in the dorsal, compared to the ventral locations. Pearson’s correlation coefficients (here and throughout, unless otherwise mentioned), are indicated above along with its p-value. **(C)** Similar to (B), for all broad spiking MEC cells (*n*=3601) combined. **(D)** Similar to (B), for all narrow spike waveform MEC cells. (*n*=945). **(E)** Similar to (B), for broad and narrow EC cells combined (*n*=5289). Red trace denotes the fraction of tuned cells in 10 equi-spaced binned of dorso-ventral location. Prevalence of tuning dropped from 80.0% to 30.3% across these 10 bins. **(F)** 3D plot depicting the anatomical location of EC units, registered to the Allen CCF atlas. Color denotes the strength of tuning. Black circles correspond to the 3 cells in (A). **(G)** Similar to (F), colored for LEC vs. MEC.

Dorso-ventral gradient in other metrics of neural selectivity such as the widths of tuned peaks, and the direction selectivity was also seen. In fact, the gradient persisted even when randomly flashed bars were used that greatly reduced neural tuning to the moving bar (see below). Thus, the gradient of tuning was a robust phenomenon, insensitive to the specific parts of the entorhinal cortex, or the neural cell type and the stimulus properties.

### Multiple mechanisms for directional encoding of the moving bar of light

To investigate the nature of visual tuning as a function of its movement direction, we considered the neural responses separately in front-to-back, i.e. naso-temporal (NT) and back-to-front, i.e. temporo-nasal (TN) directions. Given the similarity of visual tuning strength and anatomical gradient in the excitatory and inhibitory cells as well as across EC sub-layers, we pooled them as a single population for further analysis. Cells showed a variety of directional responses. Some cells had virtually identical responses in the two movement directions, having similar firing rates, strengths of tuning and the tuning curves, indicating a direction-independent representation (Fig. 3A). However, these were a minority. Many cells preferred the same location but had significantly different firing rates in two movement directions (Fig. 3B). Yet other set of cells showed significant tuning in both directions, but the tuning curves were uncorrelated (Fig. 3C), or even negatively correlated with each other (Fig. 3D). Additionally, many cells showed tuning in only one movement direction with a drop in firing in the non-tuned direction (Fig. 3E). Other cells had similar mean rates in the two movement directions but showed significant tuning in only one direction (Fig. 3F).

**Figure 3.**
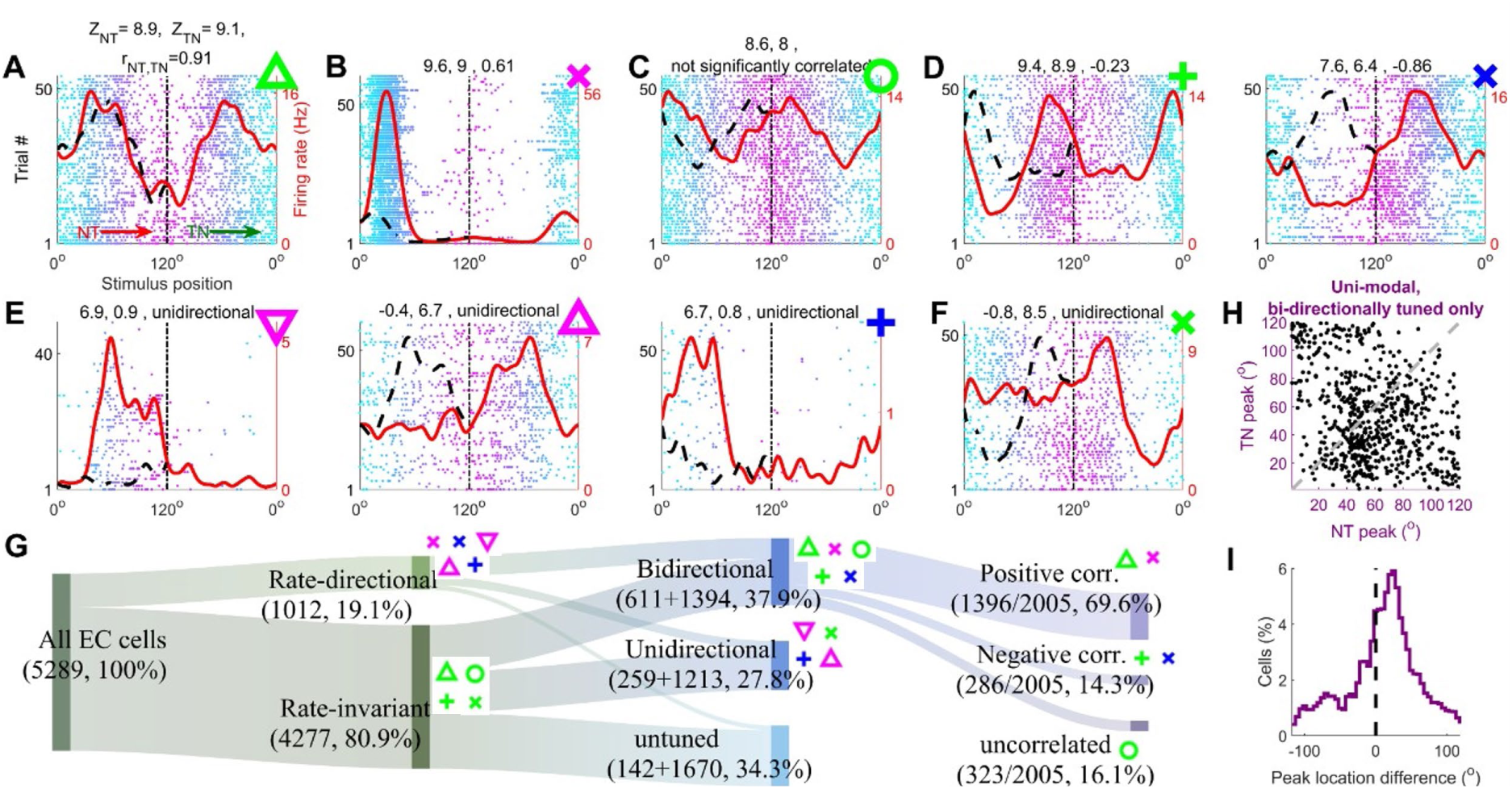
Visually evoked activity shows three types of directionality. **(A)** Example cell showing tuned responses during naso-temporal (NT) as well as temporo-nasal (TN) motion of the bar of light. Similar format as Fig. 1(D). Black dotted trace is the same response as red, but its TN response is flipped and overlaid on the NT response for comparison. Correlation between these responses is indicated above and was significant (*p*=6.3×10^−47^). Strength of tuning was computed separately for both movement directions (see *Methods*) and is also indicated above. Note-this cell had comparable mean firing rate in both directions. Cells without significant mean rate change are indicated with green symbols through (E). **(B)** An example cell with bidirectional tuning but different mean rate across two directions. Although the mean rate changed, the tuning curves were significantly positively correlated (*p*=9.6×10^−14^). **(C)** Similar to (A), for a cell with tuning in both directions that was not significantly correlated (*p*=0.17). **(D)** Similar to (A), for two cells with significant (0.01 and 1.4×10^−36^) negative correlation between tuning curves. **(E)** Three example cells with tuning as well as mean rate changes across the 2 directions, with higher firing rate in the tuned direction. **(F)** Example cell with tuning in only TN direction, with similar mean rate in both directions. **(G)** Flow chart depicting cell counts across different categories of directionalities. Tuning curve correlations were only quantified for the bidirectional cells and indicated as a percentage of total (2005) bidirectional cells. Symbols track cells from (A) through (F). **(H)** Scatter plot of peak locations, for bidirectional cells with a single peak in each direction (n=673). Note the concentration of data points to the right of the negative unity line, corresponding to a population bias between NT and TN peaks of the same cell. A random scatter of ∼1° is added for better visualization. **(I)** The difference in the peak location was significantly biased away from zero (Student’s t-test, *p*=1.3×10^−3^), with a median bias of 12 frames, corresponding to a difference of 200.0ms, equivalent to a 100ms retrospective bias for neural responses in each direction, assuming that the cells fired maximally to the stimuli at the same location in both directions but with a latency.

We used several metrics to quantify these diverse, directional responses. First, extensive work in the visual areas(37,39,64–69) has used significant differences in the mean firing rates along the two directions to quantify directionality; resulting in ∼40% directionally tuned cells in the primary visual cortex. Hence, we used the same metric (fig. S4, see *Methods*). Only 19.1% (1012/5289) of entorhinal cells significantly changed their mean firing rates between the NT and TN directions of the stimulus motion (Fig. 3G), termed rate-directional. Firing rate modulation index (see *Methods*) for the rate-directional cells was modest, averaging 0.16, and exceeded 0.33 (i.e., 2*x* change in firing rate between directions, or more) very rarely (13.1% of rate-directional cells). The majority of cells (80.9%) had no significant difference in their firing rate, hence termed rate-invariant (see *Methods*).

Instead of mean firing rates, neurons can encode the motion direction based on the tuning strengths in the two directions. While such coding scheme is rarely reported in the cortico-hippocampal circuit it was recently found in the hippocampal area dorsal CA1(70). Hence, we computed the strength of tuning separately in the two directions using z-scored sparsity as above (see *Methods*). Cells that had significant tuning in only one direction of motion were termed unidirectional (27.8%) and those that were significantly tuned in both directions were bidirectional (37.9%) (see *Methods*). Most (86.0%) rate-directional cells had significant tuning for at least one direction. The rate-invariant cells split roughly equally into untuned (1670/4277, 39.0%), unidirectional (1213/4277, 28.4%) and bidirectional tuned (1394/4277, 32.6%). For the population of unidirectional cells, the direction with significant tuning (or the direction with better tuning for bidirectional cells) did not significantly have a greater firing rate (µ=5.8Hz tuned vs. 5.6Hz untuned, fig. S4).

In addition, amongst the cells that were bidirectionally tuned, the tuning curves differed between the two movement directions. There were three types of responses: 69.6% of tuning curves were similar in both directions (i.e. significant positive correlation), 16.1% were unrelated (no significant correlation) and 14.3% had the opposite structure (significant negative correlation) (see *Methods*).

Similar to the bidirectional cells, the unidirectional cells also had an overrepresentation of significant positive correlations (56.0%), suggesting that the untuned direction response often correlated with the tuned direction response (fig. S4). Thus, not only do the majority of entorhinal neurons encode the location of a visual object, but they also encode its motion direction using three distinct mechanisms. Relative prevalence of these diverse directionalities was similar between LEC and MEC (fig. S5).

### Single peaked retrospective responses

Many medial entorhinal cells have multipeaked structure during spatial exploration(1,2,9), even on 1D tracks(14,55), and they show predictive coding(71,72). In contrast, such multipeaked structure is uncommon in the lateral entorhinal cortex(20,30). Hence, we quantified the size, number and locations of the visual receptive fields of EC cells and their response latency using a recently developed peak detection algorithm(73) (fig. S6, see *Methods*). The peaks of the visual responses were diverse in widths (mean±s.e.m.=76.5±0.9°) corresponding to a duration of ∼1.3s. Most cells had one peak per tuned direction (fig. S6). Similar patterns were observed in the medial and lateral parts in contrast to stark differences between the medial and lateral responses during spatial tasks(17,20).

The distribution of peak locations was significantly different from uniform in both the NT and TN directions, with noticeable accumulation of NT peaks around ∼50° and TN peaks around 30° (fig. S6). Similar pattern was observed at the level of single cells within the bidirectional population, with the distance between the peak locations in the NT vs. TN directions showing a consistent and significant bias (Fig. 3H,I) with a median peak lag of +12°. For stimulus moving at the speed of 60°/s and considering the same lag in both directions (+6°), this corresponds to a latency of 100ms, i.e. retrospective responses after the bar passes a certain location. This lag is quite consistent with the 138ms lag reported for the downstream brain region in the rodent CA1 under a similar conditions(70), and with the upstream LGN and V1(74–78) lags of ∼60ms in primates(75). This supports the idea of information propagation from the visual areas to entorhinal along the anatomical hierarchy(79,80). Similar to the response latencies of LEC and MEC to the parietal cortical activity under anesthesia(81,82), LEC latency was shorter than MEC in these recordings, but the difference was not significant (fig. S5). In sum, EC neurons were bidirectional more often than uni-directional, with typically a single, ∼75° wide peak in their tuned directions, and these signals seem to take about 50ms to propagate from the visual areas to the entorhinal cortex. This is in contrast to predictive coding during spatial navigation(71,72).

### Reduced but significant visual tuning when sequential motion is abolished

Visual cortical neurons encode not only the position of smoothly moving(64,65,67,78), i.e. a continuous-sequence of bars of light, but also the briefly flashed stimuli at random locations on a screen(38,75), i.e. random-sequence, indicating history-independent responses. To dissociate these two mechanisms in the entorhinal cortex, we used the same bar of light as above, but briefly (only one frame at 60Hz screen refresh, i.e. 16ms) flashed at random locations, chosen uniformly across the entire screen, i.e. random-sequence (Fig. 4A, See methods). The same random-sequence was repeated for 50-60 trials.

**Figure 4.**
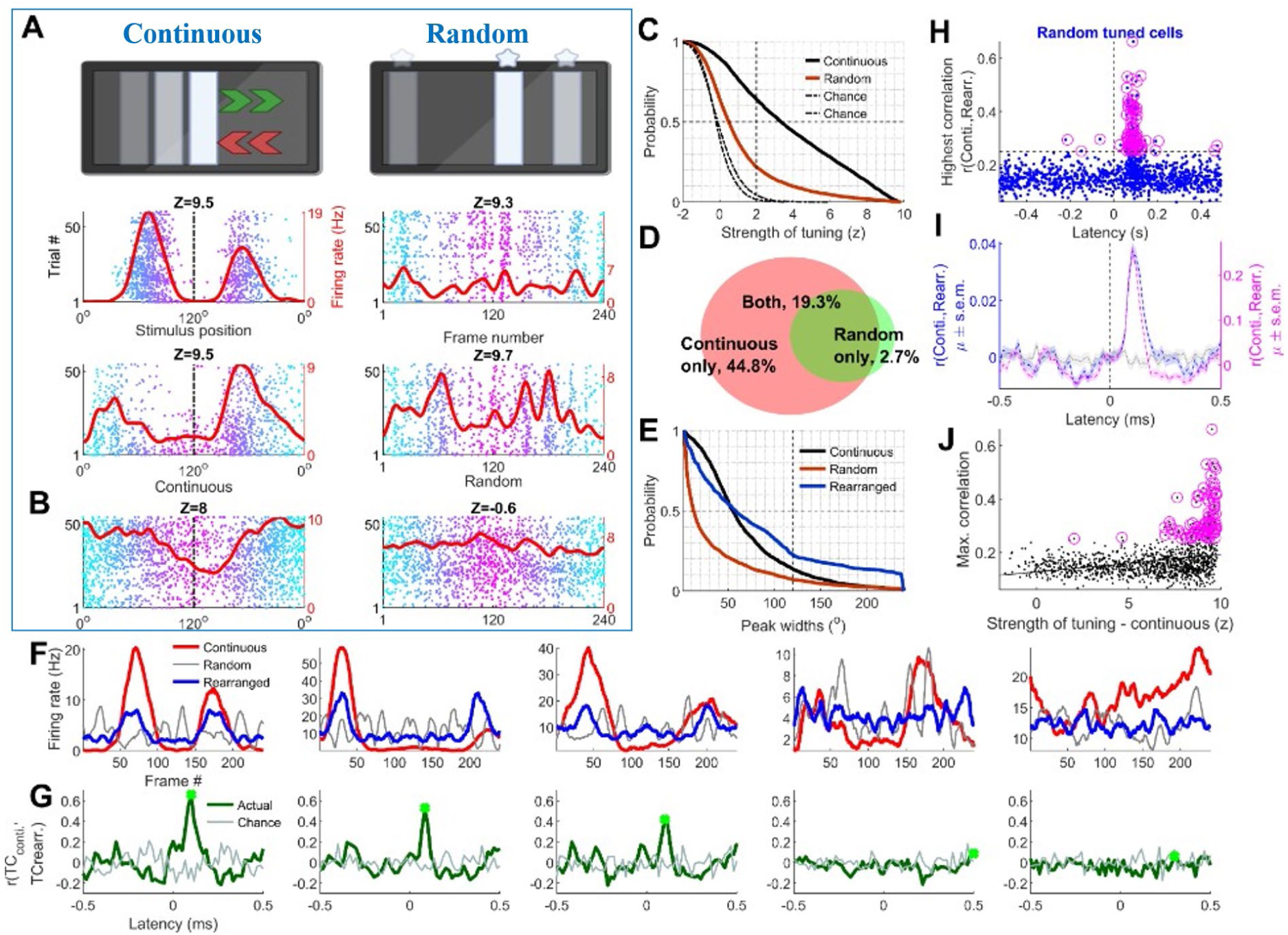
Visual tuning reduces from 69% to 22% of EC cells when sequential motion is replaced by a randomly hopping bar of light. (**A**) Top – Illustration of the experiment where the same bar of light is flashed at random locations, thus abolishing temporal coherence, called random-hopping bar. Middle and bottom panel show two example cells which remained tuned, but displayed multiple, narrow peaks for the random condition. **(B)** Example cell with tuned response to the sequential motion, but not to the random bar. **(C)** Cumulative histogram of the strength of tuning. 63.9% of cells were tuned for the sequential motion, whereas 21.9% were tuned for random hopping, i.e. 63% reduction in tuning (*p*<10^−200^). Fraction of cells tuned for the random condition was significantly greater than chance (21.9% vs. 4.4%, *p*=1.8×10^−111^). Sparsity calculations are not affected by the ordering of the bins (except for the effect of smoothing); z-scored sparsity on rearranged responses (at a fixed latency of 112.7ms) was very similar 21.5% but significantly lower than above (*p*=1.2×10^−27^) (data not shown). **(D)** Venn diagram for fraction of tuned cells. Most cells that had tuning for the random condition also had tuning for the continuous presentation of the bar of light. **(E)** Peaks widths for the random condition were narrowest (34.5°), and significantly smaller than that for the sequential bar of light (68.2°). Upon rearrangement of the random response (at optimal latency), widths were even larger on average (82.4°). All distributions were significantly different from each other (*p*<1.8×10^−15^). This analysis was restricted to 1019 cells with tuning for both the continuous and random bars of light. **(F)** Tuning of 5 example cells, compared between the continuous (red) and random bar of light stimuli, with (blue) or without (gray) rearrangement. For the first 3 cells, the red tuning curve typically has 1 or 2 strong peaks that can be recaptured by rearrangement. In comparison, the gray trace showing tuning for the random stimuli had multiple narrow peaks. By construction, this tuning curve was symmetric around the midline (120°) and did not have any directional information. First cell here corresponds to the first example from (A). **(G)** The rearrangement process was performed at multiple latencies, and the corresponding correlation between continuous and rearranged responses is shown in green. Gray trace denotes chance level correlations, using shuffled tuning curves (see *Methods*). Note-for the first 3 cells, there is a strong peak at small ∼100ms latency. This is missing in the last 2 cells. Correspondingly, in (E), these 2 cells’ reconstruction does not resemble continuous tuning curve. **(H)** Scatter plot of the maximum correlation and the latency at which that correlation was obtained. This comparison was restricted to cells with significant tuning for the random sequence (n=1163). Magenta circles denote cells for which this correlation was significant (after Bonferroni correction (see *Methods*)) and above +0.25. Note – the increased density of dots around a latency of ∼100ms. **(I)** Average (±s.e.m.) of correlation vs. latency responses for all cells (blue), and for cells with significant correlations>0.25 (magenta). The peak of this distribution for both populations is at 100ms latency. **(J)** The similarity between the sequential and the rearranged tuning curves was significantly and positively correlated with the strength of tuning for the sequential bar of light (*r*=0.29, *p*=5.2×10^−24^). (n=1163).

The mean firing rates of neurons were quite similar between the continuous versus random sequence conditions (µ=5.8 vs. 5.5Hz). 31.1% of cells had significantly higher firing during continuous-sequence, while 19.8% had higher firing during random-sequence, and the remaining 49.1% did not change mean firing rate significantly (fig. S7). Thus, the random-sequence did not cause a systemic shutdown in EC firing, and neurons responded to the random-sequence by subtle changes in mean firing rate, with a small bias for the continuous condition.

Compared to the mean rate, a clearer effect was seen for the tuning curves, which were quite different under the two conditions for virtually all cells. Some entorhinal cells spiked reliably across trials in specific segments of this random-sequence, with multiple, narrow bursts of activity (Fig. 4A) and similarly short periods of silences. However, the majority of entorhinal cells lost their tuning during the random-sequence (Fig. 4B). As a result, the prevalence of visual tuning reduced drastically from 64.1% in the continuous-sequence condition (n_total_=5289) to just 22.0% in the random-hopping condition (Fig. 4C), i.e. to a third of continuous. Conversely, virtually no neurons (2.7%) were tuned for random-sequence without the continuous-sequence tuning (Fig. 4D).

Are the random-bar induced responses related to those evoked by the continuous-sequence? Neural responses in the visual cortex appear after a short latency from the moment of stimulus onset. This latency could be variable across cells. Similar results may hold in the entorhinal cortex. Hence, we used a recently developed technique(73) (Fig. 4F, see *Methods*) to determine the cell-specific lag and rearrange the tuning curves from the random-hopping condition to resemble that during continuous condition. The correlation coefficient between the continuous and re-arranged random condition tuning curves provided an estimate of their similarity (Fig. 4G) (see *Methods*). For the vast majority (1004/1163, 86.3%) of cells, these correlations were not significant (Fig. 4H), even after considering a wide (±500ms) range of possible latencies. Successful rearrangement was only seen in cells that were tuned for NT as well as TN movement directions with similar tuning (i.e., bidirectionally tuned and positively correlated tuning curves) (fig. S7). But the converse was not true: out of all bidirectional, positively correlated tuning cells (n=1396), majority (1171, 84%) could not be successfully rearranged.

Amongst the cells with random-bar tuning, correlation was significant, i.e., rearrangement was successful for a small fraction of cells (159/1163, 13.7%), out of which half (87/159) had a correlation above 0.25; optimal latency for the reconstruction in these cells had a median of 87.7ms (Fig. 4G,H). This agrees well with the latency of 100ms obtained by comparing NT and TN tuning curves above (Fig. 3H), and in agreement with latency measurements in other studies across the visual hierarchy(74–77,79). This subset of cells had a stronger tuning in the continuous condition, compared to the rest (Fig. 4J). Taken together, entorhinal visual responses were strongly history-dependent, with a minor subset where random and continuous sequences elicited reconcilable tuning curves.

Unlike the continuous motion condition with ∼1 peaks per cell, random sequence generated multipeaked responses with ∼4.2 peaks per cell. Amongst the cells that were tuned in both conditions, the mean peak width for the random condition was ∼2*x* smaller than that for the continuous condition (34.5° vs. 68.2°, Fig. 4E) (see *Methods*). However, after the rearrangement these narrow peaks coalesced across adjacent spatial bins and the resulting peaks appeared broader than that during continuous motion (82.4° vs. 68.2°, Fig. 4E). This was true even after considering the cumulative width on a cell-by-cell basis (fig. S7). As a result, even for this subset of cells, the tuning strength was weaker for random hopping than continuous motion, further demonstrating that sequential motion generated better tuning for virtually all the cells that encoded the visual stimulus, which are the majority of entorhinal cortical cells.

## Discussion

The entorhinal cortex has been implicated in a wide range of tasks and diseases. Entorhinal neurons in the rodents have been investigated largely in the context of spatial mapping(1,2,4,6,11,29,30,58,59) and in humans in the context of visual recognition and memory(51,52,83,84). All of these involve vision, but they also involve many other sensory-motor-memory related variables resulting in a range of seemingly disparate and complex responses ranging from grid cells in rodents to concept cells in humans. Towards uncovering the basic building blocks underlying entorhinal complexity, we deployed the set of stimuli that generate robust and mechanistically explainable responses in the upstream primary visual cortex without any self-motion, reward or task. Namely, we investigated neural selectivity to a simple vertical bar of light, moving passively on a screen. Surprisingly, not only did we find highly selective responses, but this selectivity was widespread and present in a majority of neurons in both MEC and LEC. Visual tuning in many EC neurons was similar to that in the primary visual cortex, with high firing rates when the bar was in a restricted region of the screen, and virtually no spiking elsewhere. This level of prevalence is greater than many functional cell types in rodents e.g. grid cells(1,3,58), border cells(6), speed cells(8), object-vector cells(11) etc., which are found only in the medial entorhinal cortex, or time cells(16), olfactory memory cells(25,28) etc. which are found only in the lateral entorhinal cortex. While prevalence of purely visual tuning is comparable to(10), or less than(9) the fraction of spatial cells in MEC, they are greater than spatial or any functional cell type in LEC. Further, visual selectivity reported here is found in LEC as well as MEC, unlike prior work showing a major dissociation between these parts of the EC(17,20–22,29,85), based on their selectivity for space (MEC preferred space) as well as based on the differences of their afferent inputs(85–89). Moreover, while the majority of the reported entorhinal functional cell types are found only in the principal, excitatory(1,2) cells that are broad spiking, the visual responses were present in the majority of the inhibitory neurons as well (fig. S2). Thus, the purely visual tuning described here could be the unifying functional property across all of EC.

EC neurons in humans encode the turning direction in a VR environment(90), for imagery of visual images(84), as well as categories of visual images(52). Our results demonstrate that entorhinal cortex in rodents also exhibits a widespread representation of passive visual motion, similar to visual cortex(38) and hippocampus(70). Unlike hippocampal spatial selectivity that reduces considerably during immobility(91), EC visual selectivity was unaffected by running (fig. S8). Similarly, pupil dilation, a common estimate of arousal(92–96), did not have a significant effect on neural selectivity.

Visual input reaches the entorhinal cortex via a variety of pathways. Parietal cortex projects directly to the entorhinal cortex and sends projections to both postrhinal and perirhinal cortices(17,19,97). The former projects mainly to MEC(87) and the latter to LEC(87,88) (but see Ref.(23,98,99)). Other, visually responsive brain areas such as V2(80) and retrosplenial cortex project directly to the (medial) entorhinal cortex(87,100–102) and are known to receive cortical visual input(103). The postrhinal cortex has been recently shown to receive visual information from a thalamic hub conveying superior colliculus input(34), though it’s unclear if these inputs also reach the perirhinal cortex. Consistent with anatomy(103), previous study reported that MEC was responsive to drifting gratings(100). However, they did not report focused receptive fields. Other studies reported visual landmark-related activity in 10-15% of MEC cells in virtual reality(13–15). However, self-motion and reward cues too contribute to neural activity. These studies neither reported cue-locked receptive field properties, nor report any dorso-ventral gradient, nor the LEC responses. In fact, there are major differences in the anatomical projections to MEC vs. LEC and between the functional cell types during spatial exploration in these sub-regions of EC. Despite this, we found remarkably similar visually evoked responses in MEC and LEC. We hypothesize that during spatial exploration, other multisensory inputs could combine with the purely visual inputs to create these major functional differences. Such a transformation has been shown in dorsal CA1 during navigation on 1D linear tracks – distance code in VR(104,105) environments with only visual cues is replaced by absolute position code in real-world, multi-sensory tracks(106,107). Similarly, the cause of multi-peaked responses in MEC during spatial exploration(1,3) could be the non-visual, multisensory-motor cues during spatial navigation experiments since the purely visually evoked responses we report here have a single peak in MEC as well as LEC. Notably, the multi-peaked, spatially periodic grid cells are found when rats explore large, 2D mazes. These mazes can become arbitrarily large, which would yield proportionately greater grid fields(1). This is in contrast to the neural selectivity we report here, which is as a function of the stimulus position on a screen, which is finite and can be at most 360 degrees. Thus, the neural selectivity reported here is potentially related to the head-direction selective cells in MEC(4).

Medial entorhinal grid(4,58,61) and head direction(59) cells show a dorso-ventral gradient: tuning deteriorates with anatomical depth. The mechanisms are currently investigated(59–63). Similar to this, visual tuning too showed pronounced dorso-ventral gradient (Fig. 2). Significant dorso-ventral gradient was found within each mouse and within 16 out of 18 electrodes, showing that this was a robust effect (Data not shown). Such topographic organization has not been reported in LEC, or in the deeper layers of MEC(59), or in the narrow spiking inhibitory neurons in any parts. In contrast, we found such topographic organization in these parts too. Neither the mean rates of units nor the number and the location of visually evoked peaks showed such strong topography (fig. S3). But, consistent with its effect on the strength of tuning, more dorsal cells were typically more direction invariant (i.e. bidirectional with positively correlated tuning curves), with narrower peaks and with stronger tuning to the random bar of light. It is plausible that this dorso-ventral gradient reflects a preferential and prevalent connectivity of dorsal EC with the thalamo-cortical(39) and/or collicular(34) visual areas.

Visual tuning in EC showed strong dependence on movement history – many cells showed diverse stimulus motion directionality. Directionality in terms of changes in firing rate, as described typically in visual areas(37,39,64–69), was less widespread (∼20%, Fig. 3) than in many parts of the visual areas, but directionality in terms changes in strength of tuning and the location of the tuned response were more prevalent(70). Consistently, majority of EC cells lost tuning for the randomly hopping bar of light. This is unlike the primary visual cortex where briefly flashed stimuli generate substantial selectivity(38,75).

There can be many reasons for the emergence of strong sequence-dependence in the entorhinal cortex. One possibility is that the inputs from various upstream visual areas arrive in the EC at variable temporal lags, e.g. from the superior colliculus(34,89), LM and AL higher visual cortices(19), and hence they don’t summate causing the loss of tuning for 16ms long flashes. This is consistent with the observed ∼100ms response latency to the stimulus onset, which is ∼50ms greater than that reported in the visual areas. Another possibility is that the afferent visual input evokes several seconds long persistent activity(81,82) in vivo which would cause strong history dependence. This could explain not only the loss of tuning for random hopping but also the wide, single peaked visual tuning in EC. On the other hand, the EC persistent activity differs significantly between MEC and LEC whereas the visual tuning was quite similar across all parts. Thus, additional mechanisms should be investigated.

Nevertheless, even with these brief and random flashes, 22% of cells maintained significant tuning, and many cells’ random sequence responses could be reconciled with that during continuous motion. These cells had stronger tuning to the continuous moving stimulus than the rest, and were typically bidirectionally tuned (i.e., history independent). This could be because the temporal lags across inputs were less variable for these cells, causing reliable and strong response in all conditions; or because these cells received mostly visual inputs and very little multisensory inputs that could interfere with the visual tuning.

Entorhinal selectivity in rodents is predominantly measured during spatial navigation related tasks, but these tasks do not require the entorhinal cortex(108,109). The entorhinal visual selectivity described here too need not be necessary for perceiving a moving object. But the neural selectivity in this simple condition sheds light on the building blocks of neural responses during complex and demanding, spatial and non-spatial tasks. For example, the hippocampal selectivity to distance and direction in simple tasks(104,110) was strongly predictive of navigational behavior in cognitively demanding navigational tasks(111). The central finding here is that in contrast to medial entorhinal complexity and major differences across different layers and cell types, and very little selectivity in LEC, across spatial tasks, a purely passive object motion experiment drives responses in MEC as well as LEC that are comparable for tuning prevalence and dorso-ventral topography.

## Acknowledgments

We thank Dr. Massimo Scanziani and members of his lab for input and feedback. C.S.P. is an HHMI fellow of the Jane Coffin Childs Fund for Medical Research. The Neuropixels dataset in this project was obtained as part of the OpenScope program, which is operated by the Allen Institute, Neural Dynamics program and funded by the US National Institutes of Health (NIH - U24NS113646 (to Jerome A. Lecoq and Christof Koch)). We thank the OpenScope steering committee, Christof Koch and Karel Svoboda for their support. This research was also supported by the Allen Institute, founded by Jody Allen – chair and co-founder of Allen Family Philanthropies, and the late Paul G. Allen – investor, philanthropist, and co-founder of Microsoft. We gratefully acknowledge their vision and generosity, which make this work possible.

## Data availability

The datasets used herein are publicly available through the OpenScope program at https://doi.org/10.48324/dandi.000690/0.250326.0015.

## Code availability

All analyses were performed using custom-written code in MATLAB version R2020a. Codes written for analysis and visualization are available from the corresponding authors at reasonable request upon publication.

## Supplementary Materials

Materials and Methods

Author contributions

Figs. S1 to S8

References

## Materials and Methods

No statistical methods were used to predetermine sample size. The experiments were not randomized, and the investigators were not blinded to allocation during experiments and outcome assessment.

### Mice

Mice were maintained in the Allen Institute for Brain Science animal facility and used in accordance with protocols approved by the Allen Institute’s Institutional Animal Care and Use Committee. C57BL/6J wild-type mice (n = 9, 7 females), were used. See (Ref(38)) for further details.

### Surgical procedures, recovery and habituation

One month before the day of the recordings, naïve mice underwent surgery for attachment of the headframe. A pre-operative injection of dexamethasone (3.2 mg/kg, subcutaneously (s.c.)) was administered 1 h before surgery, along with Ceftriaxone (0.1mg/kg, s.c.) to reduce swelling and postoperative pain by decreasing inflammation. Mice were initially anesthetized with 5% isoflurane (1–3 min) and placed in a stereotaxic frame (Model 1900, Kopf). Isoflurane levels were maintained at 1-2% for the duration of the surgery. Body temperature was maintained at 37.5°C. Carprofen was administered for pain management (5–10 mg kg−1, s.c.) and atropine was administered to suppress bronchial secretions and regulate heart rhythm (0.02–0.05 mg mg kg−1, s.c.). An incision was made to remove skin, and the exposed skull was levelled with respect to pitch (bregma–lambda level), roll and yaw.

To enable co-registration across surgical, intrinsic signal imaging, and electrophysiology rigs, each mouse was implanted with a grade 5 titanium headframe that provides access to the brain via a cranial window and permits head fixation in a reproducible configuration. The headframe was placed on the skull and fixed in place with White C&B Metabond (Parkell), after which the craniotomy and durotomy over the entire left hemisphere is performed. A SHIELD implant, as recently described(112), was inserted in the craniotomy and covered in SORTA-clear. The implant was designed to provide access to multiple recording sites.

The mice were given 1.0-1.5mL Lactated Ringers Solution to help recover from the surgery and replace lost fluids. Surgery quality control, intrinsic imaging and habituation training as well as habituation quality control were similar to previous datasets. Health checks were performed for a week after surgery, followed by two weeks of habituation on clusters (which resembled actual recording setup), and one more week of habituation on the recording rig, before the day of recording. The habituation started with the head-fixation without any stimuli presentation; over the course of 2 weeks, longer durations of head-fixations as well as stimuli presentation (simple as well as complex, movie-like stimuli, see below) were performed.

### Probe insertion

Owing to the use of the SHIELDS implant, recording probes were inserted in the brain after removing the SORTA-clear but without additional surgical procedures. The probe cartridge was initially held approximately 30 cm above the mouse. After the mouse was secured in the headframe, the cartridge was lowered so the probe tips were approximately 2.5-mm above the brain surface. The probes were then manually lowered one by one to the brain surface until spikes were visible on the electrodes closest to the tip. After the probes penetrated the brain to a depth of around 100 μm, they were inserted automatically at a rate of 200 μm/min to avoid damage caused by rapid insertion. After the probes reached their targets, they were allowed to settle for 10mins. CM-DiI (1 mM in ethanol; Thermo Fisher, V22888) was used to localize probes *post-hoc*. Recording electrodes were assigned to the Allen Brain Atlas with a custom GUI that relied on the DiI track, electrophysiological markers (like theta power, cell/spike density) and boundaries between major cell classes (e.g., cortex vs. hippocampus).

### Data acquisition and synchronization

Methods similar to those previously described were used(38). All neural recordings were carried out with Neuropixels probes (v1.0). The electrodes closest to the tip were always used, providing a maximum of 3.84 mm of tissue coverage. Neuropixels data was acquired at 30 kHz (spike band) and 2.5 kHz (LFP band) using the Open Ephys GUI. Videos of the eye and body were acquired at 30 Hz. The angular velocity of the running wheel was recorded at the time of each stimulus frame, at approximately 60 Hz.

### Stimulus monitor

Visual stimuli were generated using custom scripts based on *PsychoPy* and were displayed using an ASUS PA248Q LCD monitor, with 1,920 × 1,200 pixels (55.7 cm wide, 60 Hz refresh rate). Stimuli were presented predominantly to the right eye, with a coverage starting at 7° to the left, and up to 113° to the right, and the monitor was positioned 15 cm from the right eye of the mouse and spanned 120° × 95° of visual space. Each monitor was gamma corrected and had a mean luminance of 50 cd/m^2^. To account for the close viewing angle of the mouse, a spherical warping was applied to all stimuli to ensure that the apparent size, speed and spatial frequency were constant across the monitor as seen from the mouse’s perspective.

### Visual motion stimuli

All animals viewed a variety of visual stimuli in two blocks of 27-30 trials each. The continuous moving visual stimulus was a vertical, white bar of light (15° in width, 95 ° height, spanning the entire vertical extent of the screen), moving at a fixed speed of 60°/sec. Each trial consisted of 4 seconds of stimuli movement. The bar of light started to move from the nasal-most location, and traversed to the temporal position, i.e. from front to back (total range 120°) in 2 seconds, and then returned from back to front in the next 2 seconds. Trials were presented without any intermittent delays. For the randomly-hopping stimulus, the same bar of light was used but instead of continuous motion it was presented in at pseudo random locations in every image frame (16.6ms). Thus, each location was flashed 2 times. The same random order was repeated for all trials and for all mice.

### Single unit categorization

Single units were classified into excitatory and inhibitory neurons using two different methods. Units with less than 0.5Hz mean firing rate across the entire recording were excluded for robust statistical estimates.

Method 1

Units with an average spike width (the duration between the minimum to subsequent maximum amplitude) greater than 0.45ms (but below 1.5ms, to prevent spurious, unphysiological spikes) were classified as broad spiking, putatively excitatory neurons. Putative inhibitory neurons were classified as narrow spiking, with spike widths below 0.45ms, similar to previous work(73,104).

Method 2

Complex spiking index (CSI), which quantifies the reduction in spike amplitude of putative pyramidal cells during burst spiking^81^ was computed for all cells (fig. S2), as done commonly for hippocampal place cells^65,76^. It was defined as

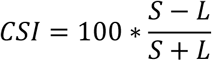

where *S* is the number of spike pairs with inter-spike interval < 20ms and amplitude of second spike less than the first; and *L* is the number of spike pairs with inter-spike interval < 20ms and amplitude of second spike was greater than the first. Units with CSI greater than 15 and a firing rate below 10Hz were also classified as putatively pyramidal (excitatory) for comparison with Method 1 above. Similarly, putative inhibitory interneurons cells were characterized to have CSI below 15 but a firing rate above 10Hz. Units with intermediate CSI were not used, to make a conservative estimate.

### Visually evoked tuning quantification by z-scored sparsity and depth of modulation

Procedures similar to those described previously were used(70,73). We binned the angular occupancy of the vertical bar and spikes in *n* = 240 bins corresponding to each image frame at 60Hz and smoothed it with a Gaussian of *σ* = 5 frames, for the calculation of the tuning curves. The strength of modulation of neural responses was quantified as sparsity *s* of the rate-map where *r_n_* is the firing rate corresponding to the *n_t_*_ℎ_ bin:

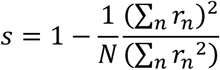

To assess the statistical significance of sparsity, we used a bootstrapping procedure(70,110), which does not assume a normal distribution. Briefly, for each cell, spike train in each trial was randomly shifted by different amounts with respect to the frame number and the sparsity of the randomized data computed. This procedure was repeated 100 times with different values of random shifts. The mean value and standard deviation of the sparsity of randomized data was used to compute the z-scored sparsity of actual data using the function *zscore* in MATLAB. The observed sparsity was considered statistically significant if the z-scored sparsity of the observed spike train was greater 2, which corresponds to *p*<0.0228 in a one tailed t-test.

This calculation of z-scored sparsity was done separately for the randomly-hopping stimuli (Fig. 4) as well as computed separately for the NT and TN movement directions of the continuous moving bar of light (Fig. 3).

Depth of modulation of firing rates was defined as (FR_max_ -FR_min_)/ FR_mean_, where FR_max_ was the highest firing rate (peak rate) across *r_n_* and similarly FR_min_ is the minimum, and FR_mean_ is the average. This metric is reported across EC sub-layers and broad vs. narrow waveform units (fig. S2).

### Statistical tests

All usages of the Kolmogorov–Smirnov (KS) test were with two-sample, two-sided hypothesis testing. All Student’s *t*-tests used were one-sample and used to test whether the distribution was centered at zero. Unless otherwise noted, correlation coefficients were computed as Pearson’s coefficients using the *corrcoef* function in MATLAB.

### Quantification of the number, location and width of visually evoked peaks

Peak detection was performed in a similar manner to that described previously(73). Briefly, the *findpeaks* function in MATLAB was used to obtain peaks with *prominence* larger than 110% (1.1*x*) the range of firing variation obtained by chance, as determined from a sample shuffled response. Each peak was assigned a score, computed as the difference between the total firing rate within that peak and total firing rate expected by chance. Peak detection and score assignment was repeated at different smoothing windows (1, 4, 16 and 64 frames, corresponding to a temporal range from 16.6ms to 1067ms) to ensure that short as well as long receptive fields were reliably detected and treated similarly. If one or more narrow peaks (putatively detected with low smoothing) overlapped with a broad one (putatively detected at higher smoothing) a comparative process was used to choose the peak(s) with a larger sum of scores. This peak comparison procedure was conducted in the increasing order of smoothing. To ensure that peaks at the beginning as well as the end of the screen were reliably detected, we circularly wrapped the tuning response, for the observed as well as shuffle data.

### Rearrangement procedure for randomly hopping stimuli responses

The randomly hopping bar of light had the same visual frames as the continuous bar of light but shuffled in a pseudo random fashion. The same random sequence was repeated for all trials and all mice. A rearrangement procedure was used to determine the extent to which neural responses in the random condition resembled that in the continuous condition. The neural response was first computed at each frame of the random sequence, keeping the frames in the chronological order of presentation. This response was then rearranged in such a way that the random sequence of frames was now in screen-based order (this was the same order as the continuous bar of light). To address the latency between stimuli presentation and its evoked neural response, which can differ across brain regions and neurons, this rearrangement calculation was repeated with variable delays between *τ* = −500 to +500 ms (i.e., −30 to +30 frames, in steps of 1 frame or 16.6ms). The correlation coefficient was computed between the neural response to the continuous stimuli and this variable delayed response at each delay as *r*_measured_(*τ*) = corrcoef(*R*_continuous_, *R*_random-rearranged_(*τ*)). *R*_continuous_ is the firing rate across frames for the continuously moving bar of light and *R*_random-rearranged_ corresponds to the random response after rearrangement, at the latency *τ*. The latency *τ* yielding the largest correlation between the two responses was designated as the putative response latency for that neuron. This latency and the resultant maximum of the correlation is reported in Fig. 4. The statistical significance of this correlation at the optimal latency was determined at a p-value of 0.05/61, to account for the multiple comparisons (61 latencies corresponding to -30 to +30 frames or -500 to +500ms).

### Firing rate modulation and tuning correlation calculations

To compare the population level changes in neural responses to stimuli properties, we computed the mean firing rate modulation index for 2 stimuli classes S1 and S2, for the same cell *A* as 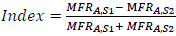 where MFR is the mean firing rate across all stimuli angles for stimuli S1. The statistical significance of this metric was assessed as follows. We mixed the trial-wise responses of the same cell across stimuli categories. Consider, for cell *A* in stimulus S1 a vector (FR_1,tr,s1_, FR_2,tr,s1_, … FR_n,tr,s1_) as the firing response across different angles, in trial *tr*. To obtain a null modulation level, the trials corresponding to stimuli S1 and S2 were mixed together and two random, non-overlapping subsets were drawn from these mixed trials. These random subsets were averaged across angles and trials to obtain the mean rates for each subset. The operation was repeated 100 times, and the modulation index for the subsets were used to z-score the actual modulation indices. Thus, a z-score of modulation index for each cell could be computed for mean rates. Fraction of cells with resultant z-scores above 2 (or below - 2) are reported in Fig 3,4 and fig. S7.

To assess the similarity of tuning curves independent of the mean rates, we computed the Pearson’s correlation coefficient between the vectors (FR_1,1_, FR_1,1_, … FR_1,n_) and (FR_2,1_, FR_2,1_, … FR_2,n_), where FR_i,j_ is the firing rate across trials for the stimuli bin “i”, for stimuli “j”. The resultant p-value accompanying the correlation coefficient was directly used to assess significance at a 0.05 level.

### Behavioral controls

To assess the contribution of arousal state on movie tuning, we re-calculated z-scored sparsity in epochs with high versus low pupil dilation. The pupil was tracked at a 30-Hz sampling rate, and the height and width of the elliptical fit was used to compute and area. For each session, the pupil area thus calculated was split into two equal halves, by using data above and below the 50th percentile (i.e., median). The resultant z-scored sparsity is reported in fig. S8. Similarly, the movement speed of the pupil was split into two equal data subsegments and compared for visual encoding in the two subsets of data. Similar to the pupil area, we also assessed the effect of animal’s running on tuning to the bar of light, by re-calculating and comparing z-scored sparsity in epochs of high (greater than median speed of the session) and low (<median) running speed.

## Author contributions

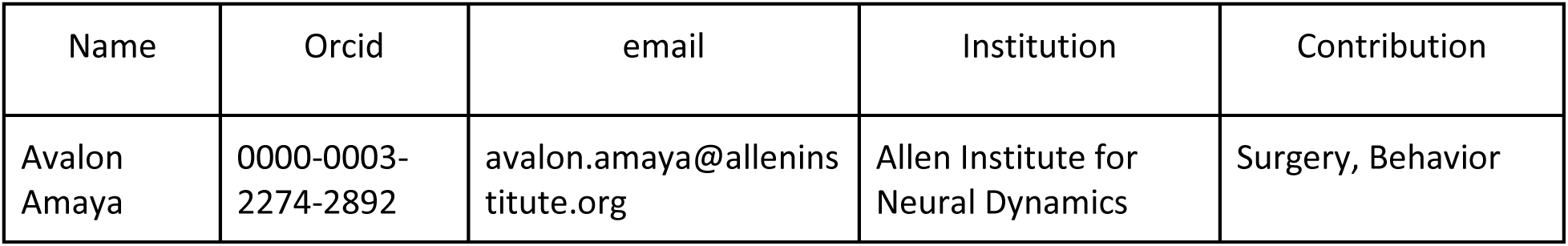
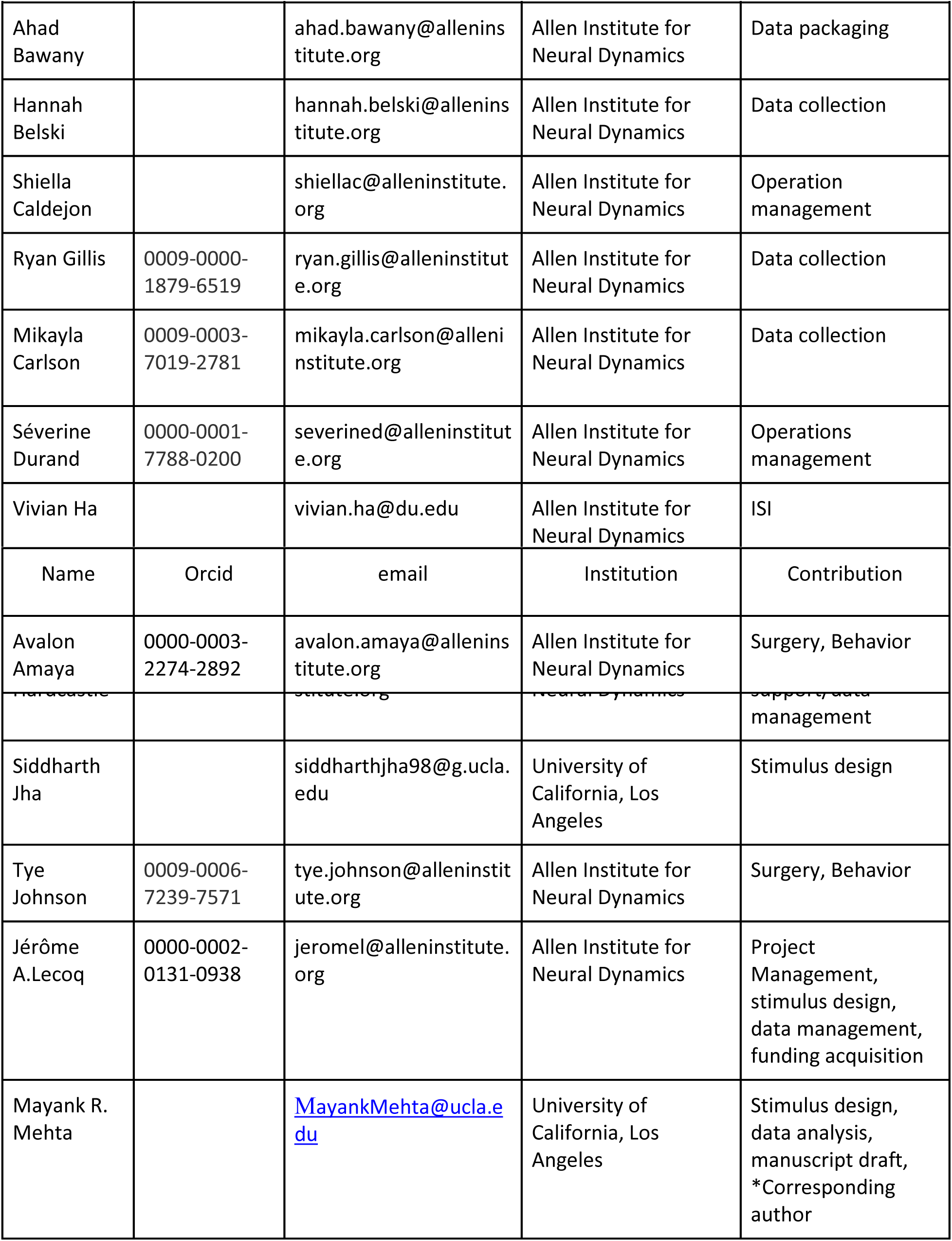
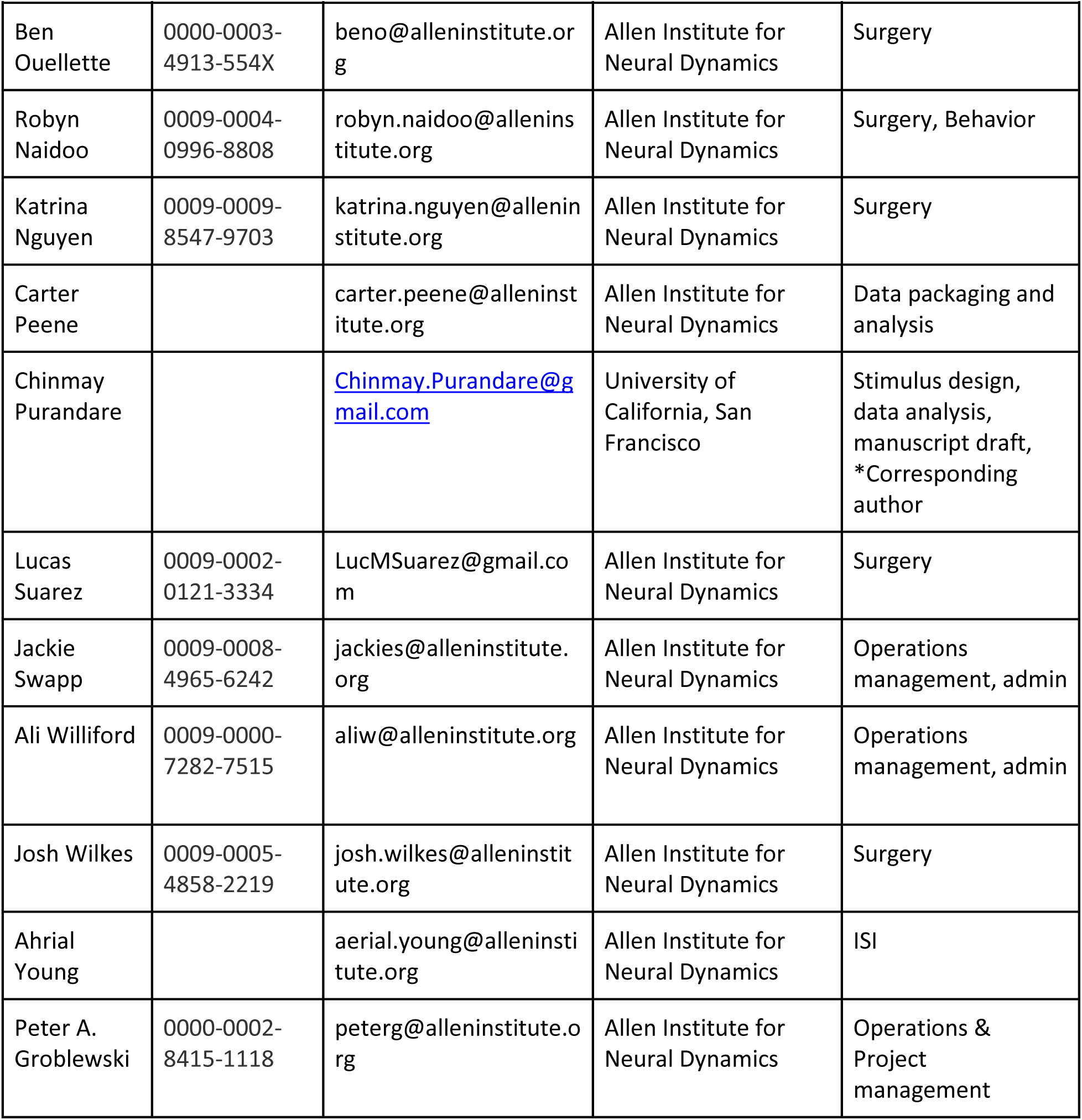

## Supplementary Figures

**Figure S1.**
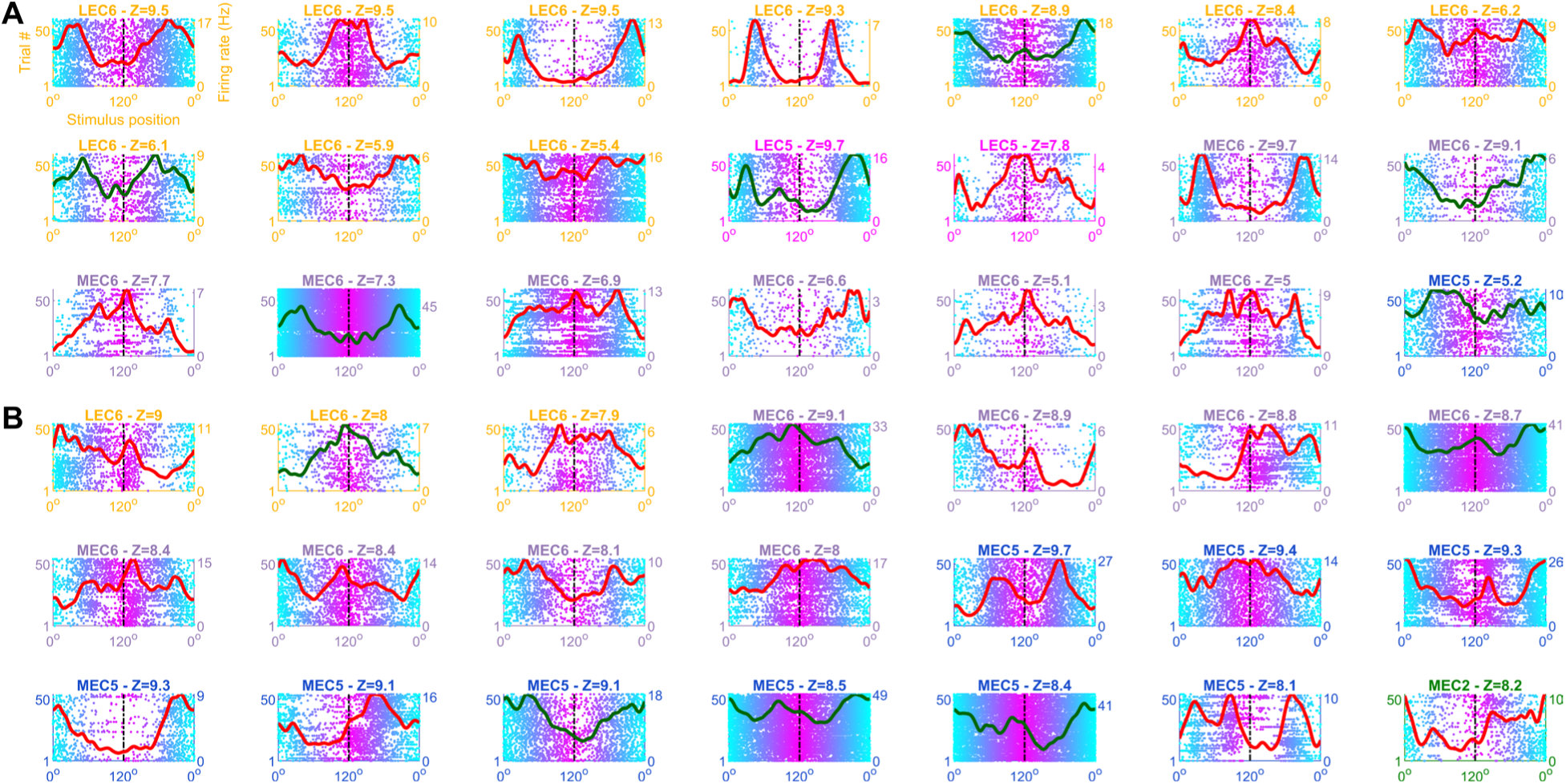
Simultaneously recorded cells show a diversity of neural responses. **(A)** 21 simultaneously recorded cells from LEC as well as MEC showing diverse tuning responses to the bar of light. Similar format as Fig. 1D. **(B)** Same as (A), from another session. Red tuning curves correspond to broad spiking cells, green traces correspond to narrow spike waveform units.

**Figure S2.**
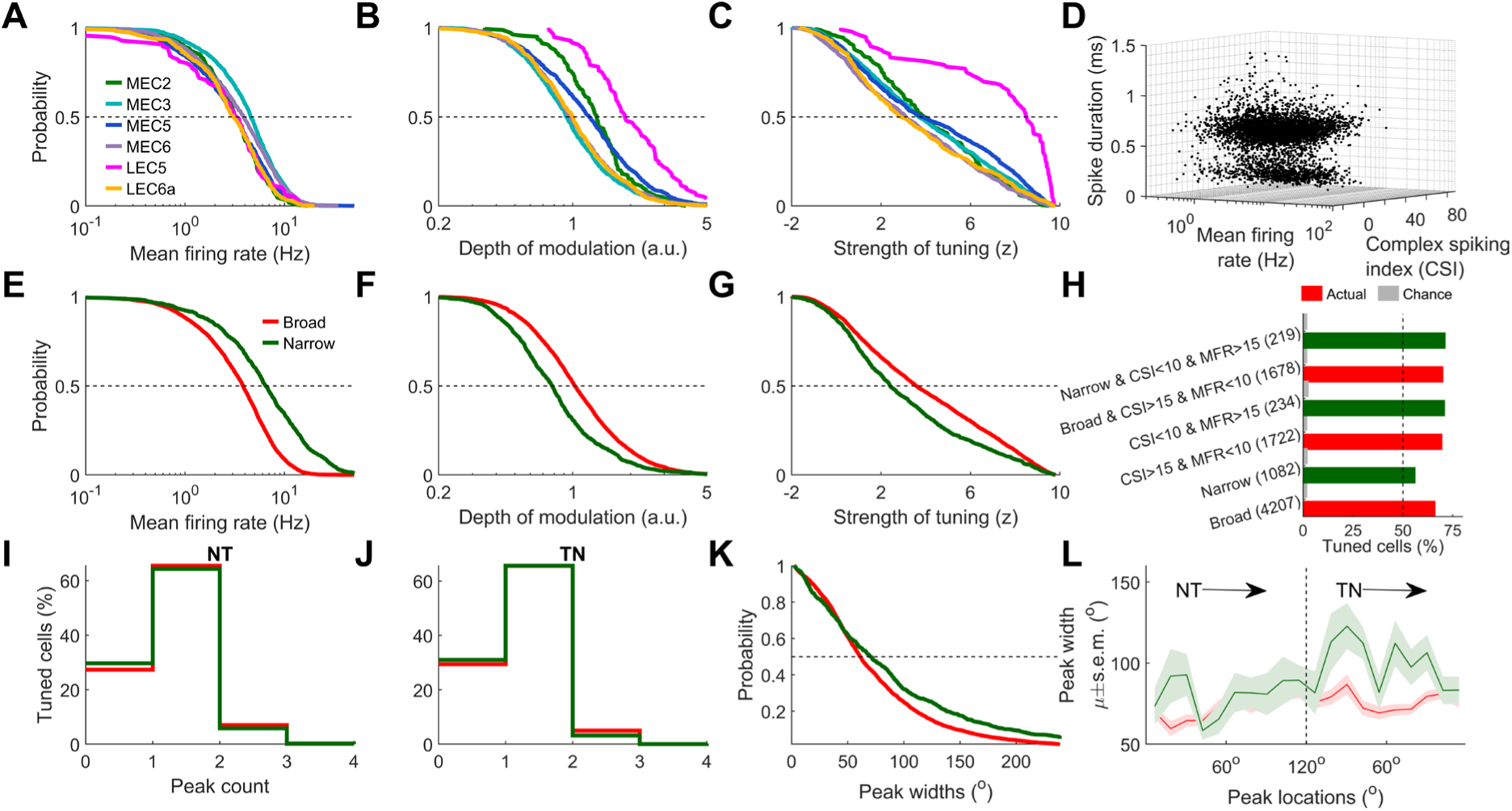
Similar visual tuning in entorhinal cortex across layers and inhibitory-excitatory cells. **(A)** Cumulative histogram of the mean firing rate of broad spiking cells from 6 layers of EC (note-x-axis is log scaled). MEC3 rates were highest (5.3±0.1Hz) and LEC6 were the lowest (3.9±0.13Hz). **(B)** Similar to (A), for the depth of modulation (see *Methods* for definition). LEC5 had the highest depth of modulation (2.3±0.12) and MEC3 had the lowest (1.1±0.02). **(C)** Similar to (A), for the strength of tuning, quantified as the z-scored sparsity (see *Methods*), was the highest for LEC5 (88.6%) and lowest for LEC6 (60.2%). To obtain a robust comparison of the activity of broad and narrow spiking units (the latter were fewer) data from all layers of both MEC and LEC were combined in panels d to L. **(D)** The 3D scatter of all EC units as a function of their mean rate, CSI and spike duration, show a clear separation in two clouds is seen along the spike duration dimension at ∼0.45ms. This was used for separating the broad and narrow spiking units (see *Methods*). **(E)** Mean firing rate of narrow spike waveform units was significantly larger than the same for broad spiking units (median=3.8Hz vs. 6.5Hz, *p*=2.3×10^−55^). **(F)** Similar to (E), comparing depth of modulation. Broad spiking units had significantly larger depth of modulation than narrow spiking units (1.0 vs. 0.78, *p*=4.0×10^−37^). **(G)** Similar to (C), the z-scored sparsity of broad spiking units was higher than that for the narrow spiking units (66.2% vs. 56.1%, *p*=1.3×10^−15^). Same data and stats as Fig. 1H. **(H)** Similar to Fig. 1H, using additional criteria of CSI to separate broad vs narrow spiking units. Imposition of CSI and mean firing rate (MFR) criteria, which can remove noisy units, resulted in a significant increase in the estimate of visual tuning prevalence (66.2% vs. 70.1% for broad *p*=2.2×10^−3^, and 56.1% vs. 71.23% for narrow, *p*=3.3×10^−5^). **(I)** Histogram of the number of peaks per cell. Most broad (65.5%) as well as narrow (64.3%) spike waveform cells were unimodal in their NT response. **(J)** Similar to (I), for TN direction responses. 65.5% and 65.6% of the responses were unimodal for broad and narrow, respectively. **(K)** Cumulative histogram of the peak widths. Broad spiking cells had significantly shorter peaks than narrow cells (74.4° vs. 86.4 °, *p*=9.9×10^−6^). **(L)** Peak width as a function of peak location for narrow and broad spiking waveforms. Peaks were significantly wider for the narrow waveform cells than the broad waveform cells, for TN (72.7° vs. 76.6°, *p*=0.03) as well as NT directions (76.3° vs. 97.2°, *p*=2.8×10^−8^).

**Figure S3.**
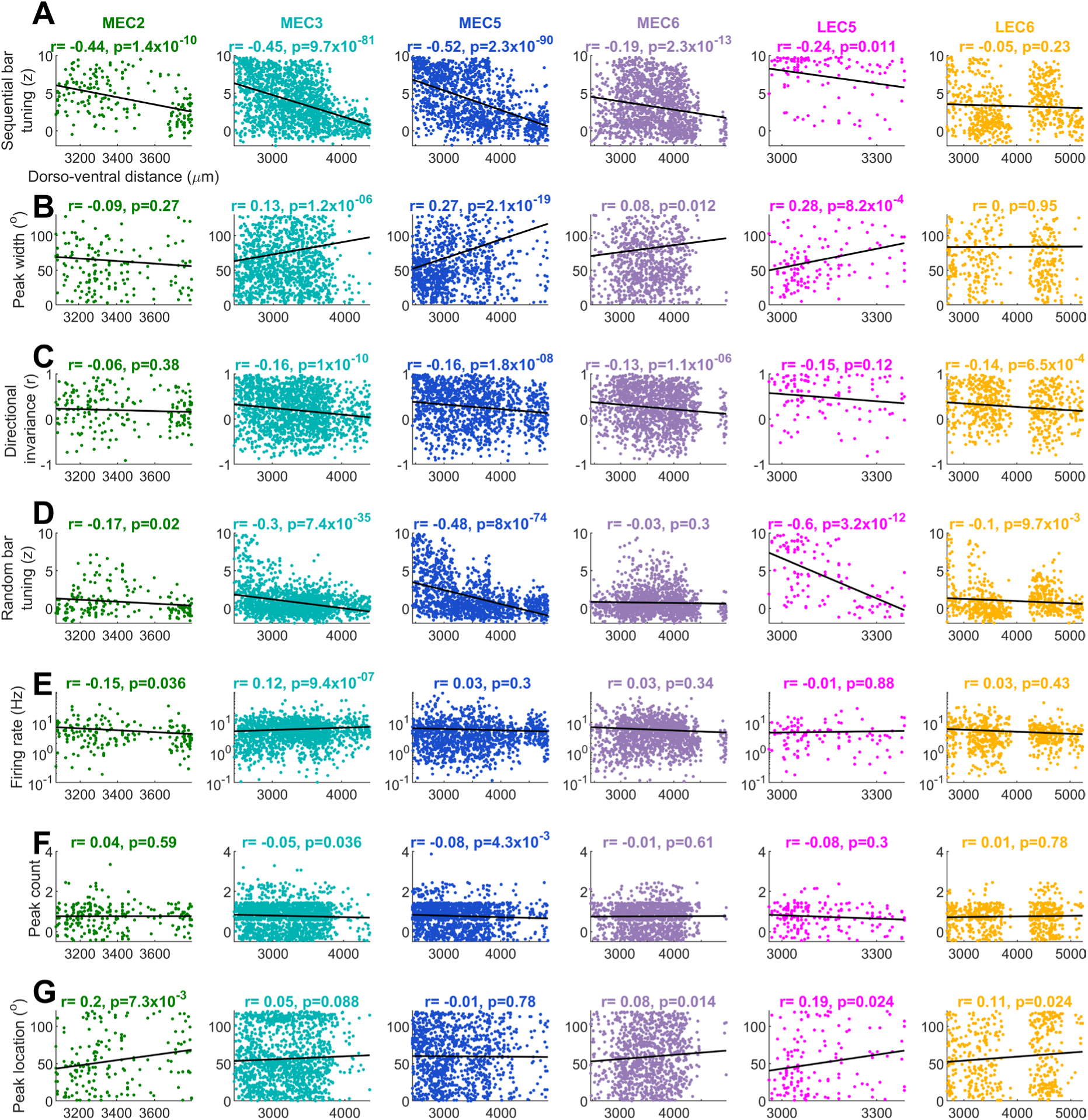
Dorsoventral dependence of neural activity and selectivity for visual tuning. **(A)** Scatter plot between the z-scored sparsity of tuning for continuous bar of light (quantifying strength of tuning) and dorso-ventral location (split by the subregion). Larger values along x-axis correspond to more ventral cells, smaller numbers correspond to dorsal (i.e., closer to the surface of the skull). Corresponding correlation coefficients and their p-values are indicated above. Only broad spiking cells are used here and through (G). Strength of tuning for the continuous moving bar of light was greater for the dorsal parts of EC than the ventral part for all sub-regions. **(B)** Similar to (A), for peak width. Dorsal neurons typically had narrower peaks, except for MEC2 and LEC6 where the relation was not significant. **(C)** Similar to (A), comparing direction-invariance, defined as the correlation between NT and TN neural responses for the same cell (unity corresponds to the cell which had the same tuning curve in both directions). Dorsal cells typically had higher direction invariance. **(D)** Similar to (A), strength of tuning for the random bar of light was significantly higher for dorsal than ventral EC neurons, except MEC6. **(E)** Similar to (A), for mean firing rate of cells. Note-y-axis is log-scaled. Mean rate did not have a consistent correlation with depth across EC layers. **(F)** Number of peaks (per tuned direction) was not significantly related to the dorso-ventral depth, except for a weak negative relation in MEC3 and 5. Only the peaks from the tuned direction(s) were used. **(G)** Similar to (A), peak location(s) for tuned cells were not related to the dorso-ventral location in MEC3 and MEC5. There was a weak relation such that dorsal neurons had a significant preference for the front, and ventral units preferred behind the animal in other sub-regions. NT and TN directions were treated separately; cells with multi-peaked responses resulted in repeated counting of their dorso-ventral locations

**Figure S4.**
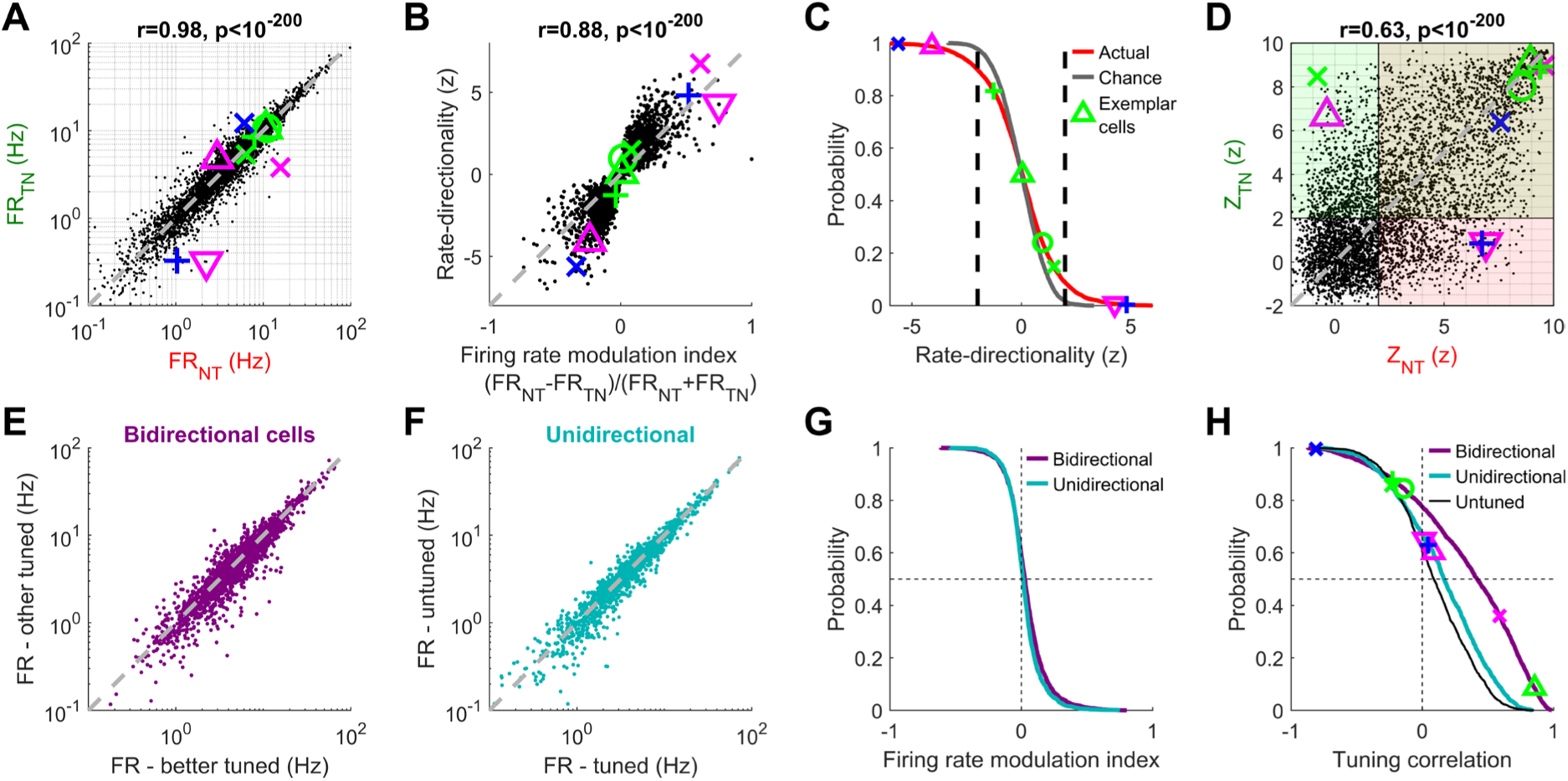
Bidirectional as well as unidirectional cells had higher firing rates in the better tuned direction, but this effect was modest. **(A)** Scatter plot of firing rate for the same cell in the NT vs TN directions. Example cells from Fig. 3A-F can be tracked here, with their symbols. 70.6% of cells had firing rate ratio within 0.8-1.25*x.* (B) As expected, the commonly used metric of rate change between NT vs. TN direction, the firing rate modulation index, was strongly correlated with the rate-directionality, quantified as the z-scored modulation index (see *Methods*). (C) Cumulative histogram of z-scored rate-directionality, for actual data and shuffles. The example green cells had <±2 zscores, indicating chance-level changes in the mean rates across two directions. (D) Scatter plot of the strength of tuning in two directions, computed as z-scored sparsity. Overlaid green and red boxes correspond to NT and TN tuned cells, respectively. (E) Mean firing rate of bidirectionally tuned EC cells (n=2010) was slightly higher in the direction with better tuning (greater z-scored sparsity) but this effect was not significant (6.7Hz vs. 6.4Hz, *p*=0.05). **(F)** Similar to (E), comparing the unidirectional cells (n=1479) for the mean rates in the tuned vs. untuned direction. Across the population, the mean rates were not significantly different (5.7Hz vs. 5.6Hz, *p*=0.64). **(G)** The firing rate modulation index between the two movement directions (Better/Other for bidirectional cells, Tuned/Untuned for unidirectional cells) was significantly different between the two populations but the effect was modest. Bidirectional cells’ ratios were significant but marginally greater (1.15 vs. 1.10, *p*=2.2×10^−5^). **(H)** Cumulative histogram of the correlation between tuning responses in the 2 directions for cells belonging to bi-vs. uni-directional categories. Bi-directional correlations were significantly greater than those for uni-directional cells (*p*=5.9×10^−62^). Unidirectional cells also had significantly greater (9.4×10^−9^) correlation than untuned cells (solid black trace), suggesting that the untuned response was still related to the tuned one.

**Figure S5:**
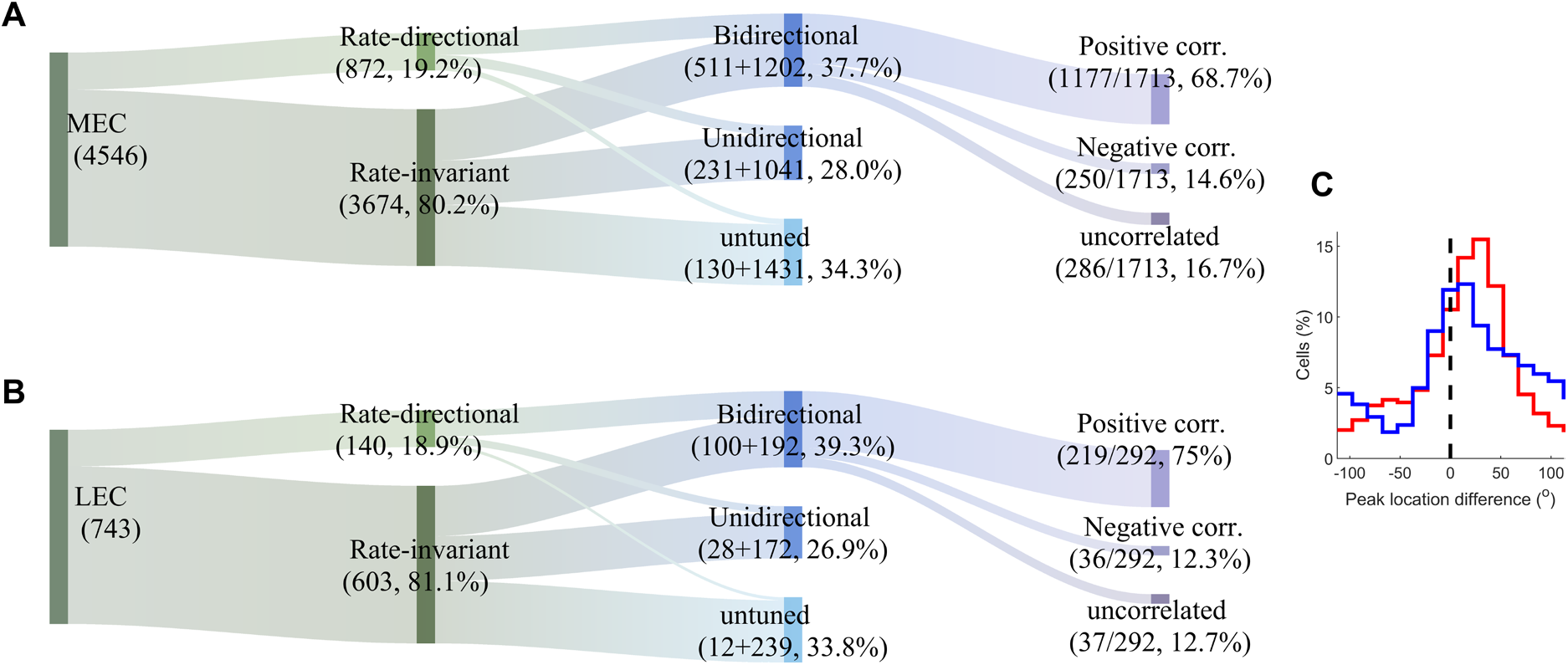
Similarity of directionality measures between LEC versus MEC |. Flow chart, similar to Fig. 3G, for **(A)** MEC and **(B)** LEC separately**. (C)** The distribution of peak location difference was not significantly different between LEC and MEC (*p*=0.1), but LEC median was lower (5°) than MEC (12°).

**Figure S6.**
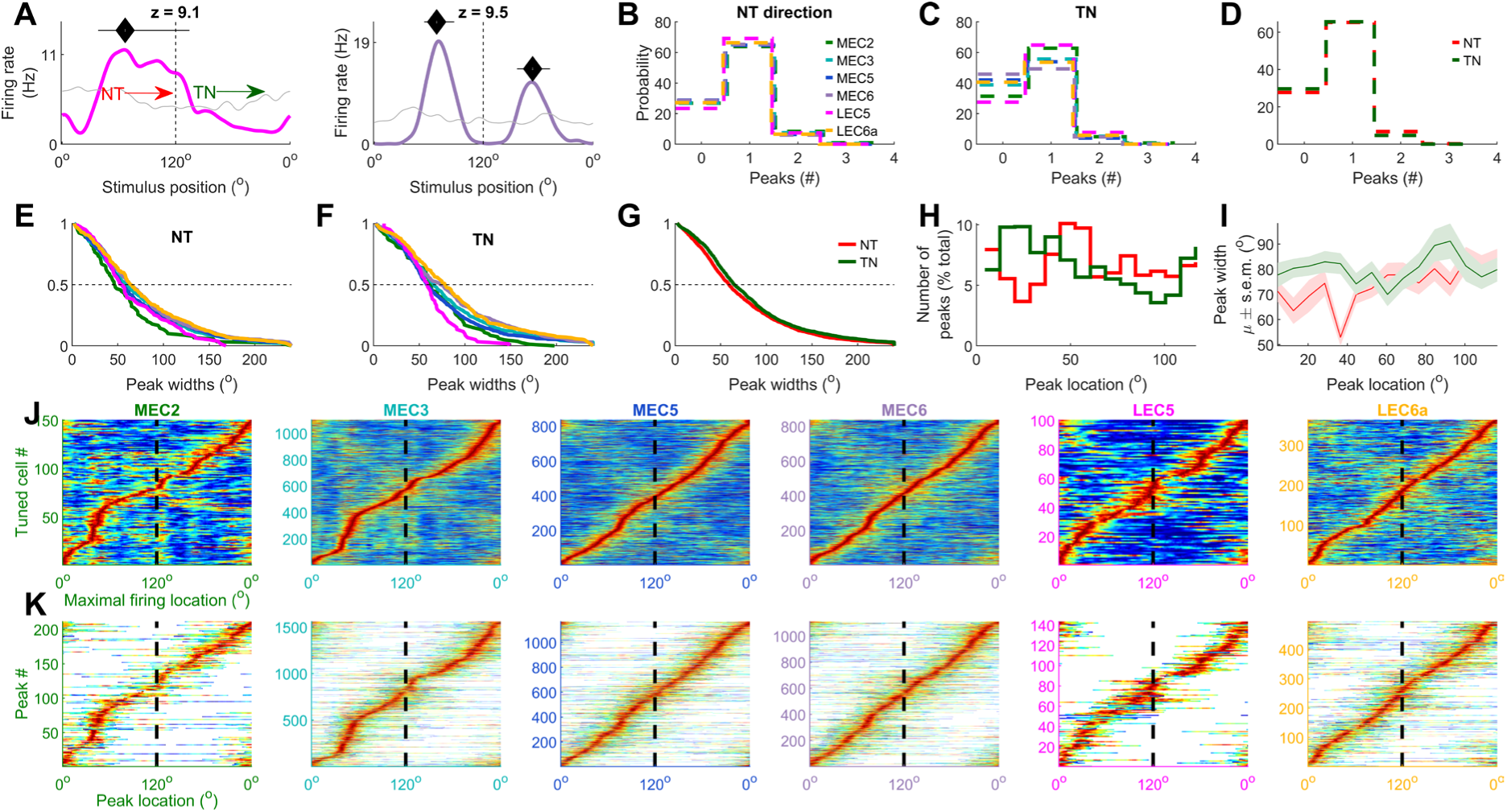
Visually evoked entorhinal cells represent all stimulus locations and are typically monotonic. **(A)** The tuning curves (colored lines) of two example cells, one from LEC5 and another from MEC6, show far greater modulation compared to chance (obtained by random shuffle of trials, gray). Black diamonds indicate peak locations (see *Methods*) and black horizontal lines indicate the peak widths of. The tuning strength (z-scored sparsity) is indicated above. **(B)** Histogram of the number of peaks in the NT direction of motion showed that most cells had only one peak, in all layers. Only cells with significant tuning in the NT direction are used here (see *Methods*). **(C)** Similar to (B), for TN tuned cells. 29.7% cells did not have a peak in NT direction, and 27.8% did not have a peak in the TN direction, even though that response was deemed selective, putatively due to the strict peak detection criteria (see *Methods*). **(D)** Similar to (B), combining EC layers, and comparing NT and TN distributions. These were not significantly different (*p*=0.55). **(E)** Cumulative histogram of peak widths in the NT direction from tuned cells. Peaks of LEC6 were widest (mean=78.7 frames), and MEC2 were the narrowest (59.4 frames). **(F)** Similar to (E), for TN direction. Peaks of LEC6 were the widest (mean=89.2 frames), and LEC5 were the narrowest (61.4 frames). **(G)** Similar to (F), combining across layers. NT peaks were significantly narrower than TN peaks (73.3° vs. 79.6°, *p*=1.2×10^−5^). **(H)** Histogram of the fraction of peaks as a function of their location. Peaks in the NT direction over-represented the locations around 50°, whereas those in the TN direction did so around 30°. This further indicates retrospective coding at the neural ensemble level (Fig. 3H,I). **(I)** Mean peak width as a function of peak location. NT peaks increased in size from the front to the back (from 71.2° to 83.7°). This effect was weaker for TN peaks (78.1° vs. 79.4°). **(J)** Stack plot of all tuned cells for a given brain region, arranged according to the location of their peak firing rate. Maximal firing rate bins span all frames, with MEC2&3 over-representing ∼50°, corresponding roughly to the center of the visual field, but only in the NT direction of motion and not the TN direction. **(K)** Similar to (J), stacking the peaks across tuned cells per sub-layer of EC, arranged based on the peak location. Color indicates firing rate in the peak (normalized between 0-1 across different peaks). Unlike (J), this plot allows for a single cell to contribute to multiple peaks.

**Figure S7.**
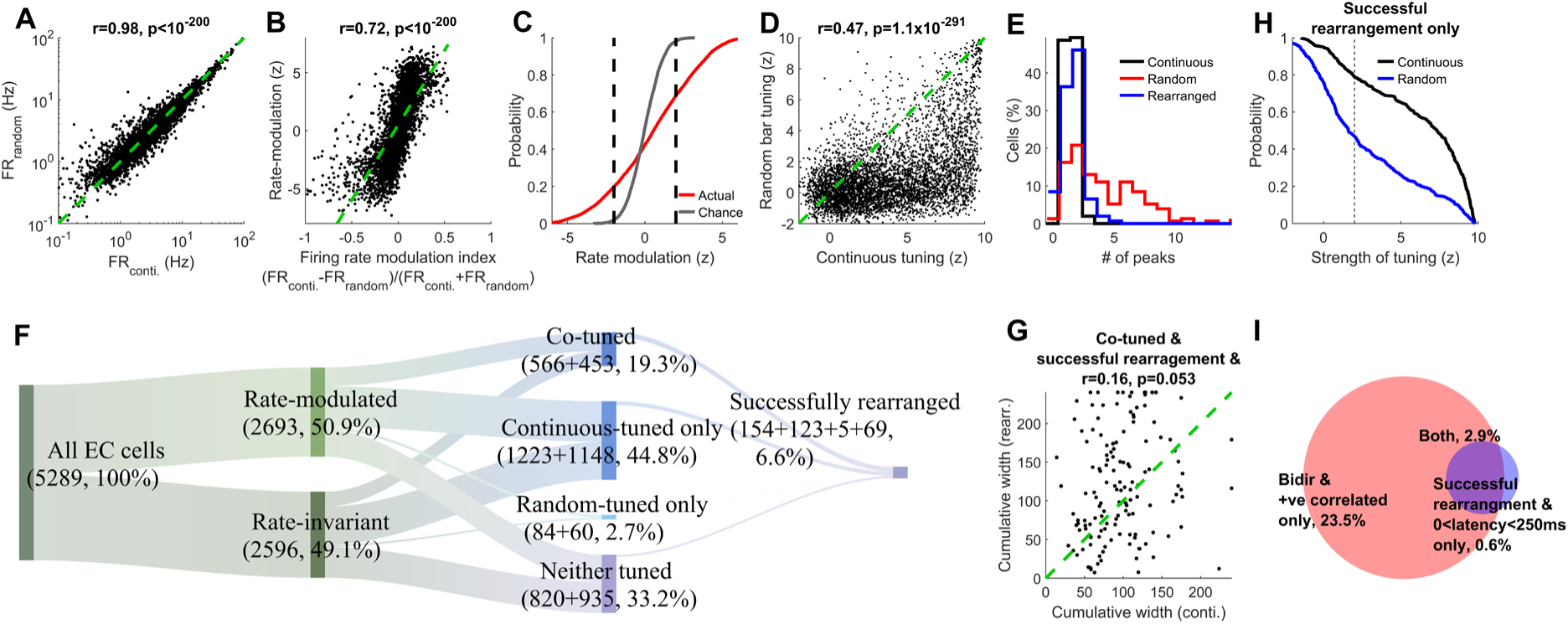
Rearrangement of random-sequence responses is successful in capturing the continuous motion responses primarily for the highly tuned bidirectional cells. (**A**) The mean firing rates during continuous and random presentations were highly correlated (Pearson’s correlation coefficient and its p-value indicated above), with a small but significantly higher firing during continuous bar (5.8 vs. 5.5Hz, *p*=0.04). 69.0% of cells had firing rates within 0.8-1.25*x* of each other for the two conditions. (B) Firing rate modulation index was computed for rate change between continuous and random conditions, similar to fig. S4B, using two different methods, the rate-modulation index and z-scored modulation. As expected, the two were strongly correlated (see *Methods*). (C) The z-scored index of the rate modulation (red) was significantly different than chance level (gray). For 31.1% of cells this difference was significantly positive (i.e. FR_continuous_ > FR_random_), 19.7% had negative, and 49.2% showed no significant difference (at a z=±2 threshold). As expected, the same for chance levels (shuffled data) was 96.2%. (D) Strength of tuning was strongly correlated between the continuous and random bar conditions but significantly larger (KS-test, *p*<10^−200^) for the continuous motion. (E) Histogram for the number of peaks for 154 cells with successful rearrangement and tuning in both conditions. Continuous and rearranged tuning curves had similar number of peaks on average (1.5 vs. 1.6, *p*=0.62). In comparison, the random (i.e., before rearrangement) condition had significantly larger number of peaks (µ=4.2, *p*<1.7×10^−19^). (F) Flow chart depicting the cell count for rate modulation and tuning across the two conditions. Roughly equal number of cells were rate-modulated as rate-invariant, and comparable fractions out of both were co-tuned, i.e., tuned for continuous as well as random bar of light (21% or 566/2702 and 17.3% or 453/2615). (G) Cumulative width, summed across all peaks for co-tuned cells were significantly positively correlated between continuous and rearranged conditions. (H) Out of the 351 cells for which rearrangement was successful (see *Methods*), 78.8% were tuned for continuous condition, and 46.7% were tuned for rearranged. This difference was significant (*p*=2.3×10^−25^). (I) Venn diagram depicting history independence in two different ways – either based on the tuning for NT and TN directions (bidirectional cells with positive correlation) or based on the cells whose rearrangement was successful at a reasonable latency (0-250ms). Almost all cells with successful rearrangement were bidirectionally tuned with significant positive correlation between their NT and TN tuning curves.

**Figure S8.**
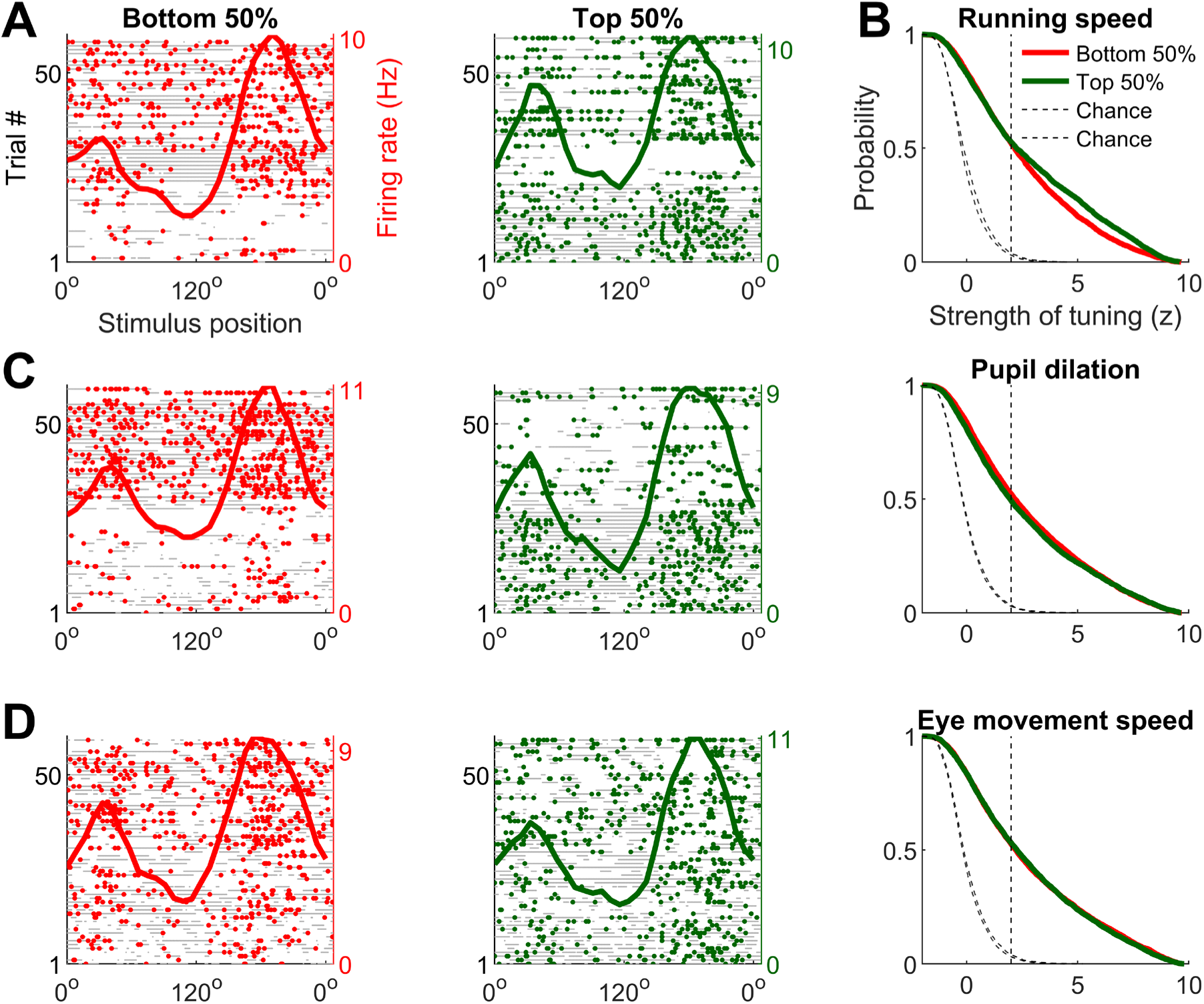
Visual tuning in EC is not affected by behavioral measures of running, arousal (pupil dilation) or eye movements. **(A)** Response of a representative EC cell was quite similar at low speeds (bottom 50%, brown panel) and high speeds (upper 50%, blue panel) (see *Methods*). Gray dots indicate occupancy (16.6ms resolution), brown (and blue) dots represent spikes. Solid traces are the average tuning curve. **(B)** Cumulative histogram of the strength of tuning for the lower and upper half of data based on running speeds. High speed data had significant but minimally greater tuning than low speed data (52.5% vs. 52.1%, *p*=4.7×10^−15^). Fraction tuned here are lower than those reported in Fig. 1 due to the reduction in the amount of data, due to splitting the entire experiment into two halves. **(C)** Similar to (A) but split based on the median pupil dilation. High pupil dilation data had significantly but only slightly greater tuning than low (52.2% vs. 49.2%, *p*=0.001). **(D)** Similar to (A) but split based on the eye movement speed. High eye movement data was not significantly different than low (52.6% vs. 53.0%, *p*=0.81).

